# Distinct dynamics and intrinsic properties in ventral tegmental area populations mediate reward association and motivation

**DOI:** 10.1101/2024.02.05.578997

**Authors:** Jordan E Elum, Eric R Szelenyi, Barbara Juarez, Alexandria D Murry, Grigory Loginov, Catalina A Zamorano, Pan Gao, Ginny Wu, Scott Ng-Evans, Xiangmin Xu, Sam A Golden, Larry S Zweifel

**Affiliations:** Graduate Program in Neuroscience, University of Washington, Seattle, WA, USA; Department of Biological Structure, University of Washington, Seattle, WA, USA; Department of Neurobiology, University of Maryland, Baltimore, MD, USA; Center for Integrative Brain Research, Seattle Children’s Research Institute, Seattle, WA; Department of Pharmacology, University of Washington, Seattle, WA, USA; Department of Anatomy and Neurobiology, School of Medicine, University of California, Irvine, CA, USA; Department of Psychiatry and Behavioral Sciences, University of Washington, Seattle, WA, USA; University of Washington Center of Excellence in Neurobiology of Addiction, Pain, and Emotion (NAPE), Seattle, WA, USA

## Abstract

Ventral tegmental area (VTA) dopamine neurons regulate reward-related associative learning and reward-driven motivated behaviors, but how these processes are coordinated by distinct VTA neuronal subpopulations remains unresolved. Here we examine the neural correlates of reward-related prediction-error, action, cue, and outcome encoding as well as effort exertion and reward anticipation during reward-seeking behaviors. We compare the contribution of two primarily dopaminergic and largely non-overlapping VTA subpopulations, all VTA dopamine neurons, and VTA GABAergic neurons of the mouse midbrain to these processes. The dopamine subpopulation that projects to the nucleus accumbens (NAc) core preferentially encodes prediction-error and reward-predictive cues. In contrast, the dopamine subpopulation that projects to the NAc shell preferentially encodes goal-directed actions and reflects relative reward anticipation. VTA GABA neuron activity strongly contrasts VTA dopamine population activity and preferentially encodes reward outcome and retrieval. Electrophysiology, targeted optogenetics, and whole-brain input mapping reveal heterogeneity among VTA dopamine subpopulations. Our results demonstrate that VTA subpopulations carry distinct reward-related learning and motivation signals and reveal a striking pattern of functional heterogeneity among projection-defined VTA dopamine neuron populations.

## Introduction

Classical models of learning and motivational processes implicate a role for mesolimbic dopamine in multiple facets of reinforcement learning and motivation. Phasic dopamine neuron activity encodes prediction-error signals which support value-based learning processes (Schultz et al., 1997; O’Doherty et al., 2017). Moreover, mesolimbic dopamine provides an incentive salience signal to modulate the strength and persistence of motivated responding (Berridge and Robinson, 1998; Bromberg-Martin et al., 2010; Niv et al., 2007; Salamone, 1999). Beyond this, when animals behave in complex tasks and naturalistic environments dopamine neurons display heterogeneous and multiplexed responses (Engelhard et al., 2019). In addition to encoding of prediction-error and incentive salience signals, dopamine neurons have been implicated in diverse processes including unconstrained, self-motivated spontaneous behavior (Markowitz et al., 2023), exploratory behavior (Harris et al., 2022), decision-making (O’Doherty et al., 2017; Cox and Witten, 2019; Saddoris et al., 2015), working memory (Adcock et al., 2006; Choi et al., 2020), sleep-wake behaviors (Eban-Rothschild et al., 2016), and social interaction (Gunaydin et al., 2014, Torquet et al., 2018; Solié et al., 2022). Further, dopamine neuron activity can encode movement and accuracy behavioral variables (Engelhard et al., 2019; Bakhurin et al., 2023), track ingestive information (Grove et al., 2022), represent information about salient and noxious stimuli (Horvitz, 2000; Brischoux et al., 2009; Lammel et al., 2011), and mediate fear association (Jo et al., 2018) and fear extinction (Salinas-Hernández et al., 2018; Cai et al., 2020).

Consistent with this pattern of heterogeneity in regulating diverse behavioral functions, VTA dopamine neurons are heterogeneous in their afferent and efferent connectivity and intrinsic neurophysiological properties (Morales and Margolis, 2017; Lammel et al., 2012; Poulin et al., 2018). To better understand how heterogeneity in the mesolimbic dopamine system contributes to motivated behavior, recent studies have emphasized the role of local control of dopamine release via postsynaptic mechanisms in the striatum for the motivational functions of VTA dopamine (Mohebi et al., 2020). Others focused on projection-specific dopamine populations and demonstrated that different regions of the striatum receive distinct dopamine signals (Cox and Witten, 2019; O’Doherty et al., 2004; Balleine et al., 2007; van Elzelingen et al., 2022), and that distinct mesolimbic dopamine pathways encode value and prediction-error information (de Jong et al., 2024). Recent work has identified differential functional roles for dopamine projections to the NAc core and NAc shell in mediating Pavlovian association (Saunders et al., 2018; Heymann et al., 2020) and appetitive and aversive motivation, respectively (Heymann et al., 2020; de Jong et al., 2019). These findings and others have been integrated into an updated model in which, rather than uniformly reflecting homogenous teaching and motivation signals, heterogeneous midbrain dopamine neurons have nuanced roles in reward-related learning and motivated behaviors (Collins and Saunders, 2020). Previous results have established differential patterns of necessity and sufficiency of NAc core and NAc shell dopamine projection populations in reward-related associative learning and motivated responding (Saunders et al., 2018; Heymann et al., 2020). However, the functional role of their endogenous activity dynamics during behavior and intrinsic neurophysiological and circuit properties remains unclear.

The VTA contains multiple types of neurons including dopaminergic, GABAergic, glutamatergic, and combinatorial populations many of which co-release neurotransmitters and neuropeptides (Morales and Margolis, 2017, Nair-Roberts et al., 2008; Parker et al., 2019). Importantly, VTA GABA neurons comprise roughly one-third of all VTA neurons (Nair-Roberts et al., 2008), regulate VTA dopamine neuron excitability through direct inhibition (Jhou et al., 2009; Johnson et al., 1992; Tan et al., 2012; Van Zessen et al., 2012), mediate ongoing motivated behavior (Van Zessen 2012), and modulate VTA dopamine neuron prediction-error responses (Eshel et al., 2015). Monosynaptic tracing studies have revealed that projection-defined VTA dopamine neurons receive inputs from a diverse array of brain regions (Watabe-Uchida et al., 2012; Tian et al., 2016, Beier et al., 2015; Lammel et al., 2012) many of which carry mixed information related to reward prediction (Tian et al., 2016). Among the numerous inputs to VTA dopamine neurons, many are inhibitory and synapse onto VTA GABAergic neurons (Soden et al., 2020), resulting in disinhibition of VTA dopamine neurons (Johnson et al., 1992; Jhou et al., 2009; Nieh et al., 2016; Yang et al., 2018). Although VTA GABA neurons have been implicated in motivated behavior, integrating these findings with an updated understanding of heterogeneous projection-defined dopamine populations has presented a challenge.

In addition to heterogeneous afferent and efferent connectivity, distinct dopamine projection populations have distinct electrophysiological properties (Lammel et al., 2008; Lammel et al., 2011; Poulin et al., 2018; Heymann et al., 2020), and ion channel expression patterns (Juarez et al., 2023; Simon et al., 2023), factors that likely contribute to their distinct functional properties. However, it remains unclear how information about reward-related stimuli and motivation during reward-seeking behaviors is encoded across specific VTA subpopulations and whether distinct intrinsic neurophysiological and circuit connectivity properties define these populations. Here we use fiber photometry and optogenetics during behavior to examine the neural correlates and functional significance of VTA dopamine subpopulations and the VTA GABAergic population in reward-related associative learning and reward-driven motivational processes. Furthermore, we use electrophysiology, targeted optogenetics, and whole-brain input mapping to assess heterogeneity among these VTA subpopulations.

## Results

### VTA subpopulations display distinct response profiles during instrumental conditioning

Subpopulations of dopamine neurons that differentially regulate reward association and motivation can be isolated in mice using genetic methods (Heymann et al., 2020). Using this approach, we sought to resolve how and when these VTA dopaminergic subpopulations encode task-related features during an appetitive instrumental conditioning task. To achieve this, we monitored neural activity in subpopulations of dopamine neurons that have been shown to differentially innervate the NAc core (*Crhr1*_VTA_ cells) or NAc shell (*Cck*_VTA_ cells) during a cued reinstatement paradigm (**Figure 1A-1B**; Nugent et al., 2017; Soden et al., 2022). We also monitored neural activity dynamics in VTA dopamine neurons as a whole (DAT_VTA_) and in VTA GABAergic neurons (Vgat_VTA_). *Slc6a3*(DAT)*-Cre, Cck*-Cre, *Crhr1*-Cre, and *Slc32a1*(Vgat)-Cre mice were injected with an AAV expressing a Cre-dependent GCaMP6m in the VTA and implanted with an optical fiber above the VTA for fiber photometry recording of time-varying bulk GCaMP fluorescence (**Figure 1D-1O**; **Figure S1A**). During acquisition sessions, a trial is initiated with an active lever-press (trial-initiation press). After a 3-s delay, the chamber house light is turned off and a compound tone-light stimulus (CS) is presented for 3-s to indicate an upcoming sucrose pellet reward. Following a 12.5-s intertrial interval (ITI) the chamber house light is illuminated indicating the availability of a new trial (**Figure 1B, left**). Lever-presses during the 3-s delay period, 3-s CS-presentation, and 12.5-s ITI are unrewarded. After acquisition mice underwent extinction sessions, during which no CS presentations or sucrose rewards were delivered (**Figure 1B, middle**). During a single reinstatement session, five non-contingent CS-presentations are delivered during a 10-min presession period. Immediately following the presession, both levers are extended and responses on the active lever leads to cue presentation following a 3-s delay but not sucrose reward delivery (**Figure 1B, right**). Mice increased their responding on the active lever during acquisition, decreased responding during extinction, and increased responding during reinstatement (**Figure 1B**).

**Figure 1.**
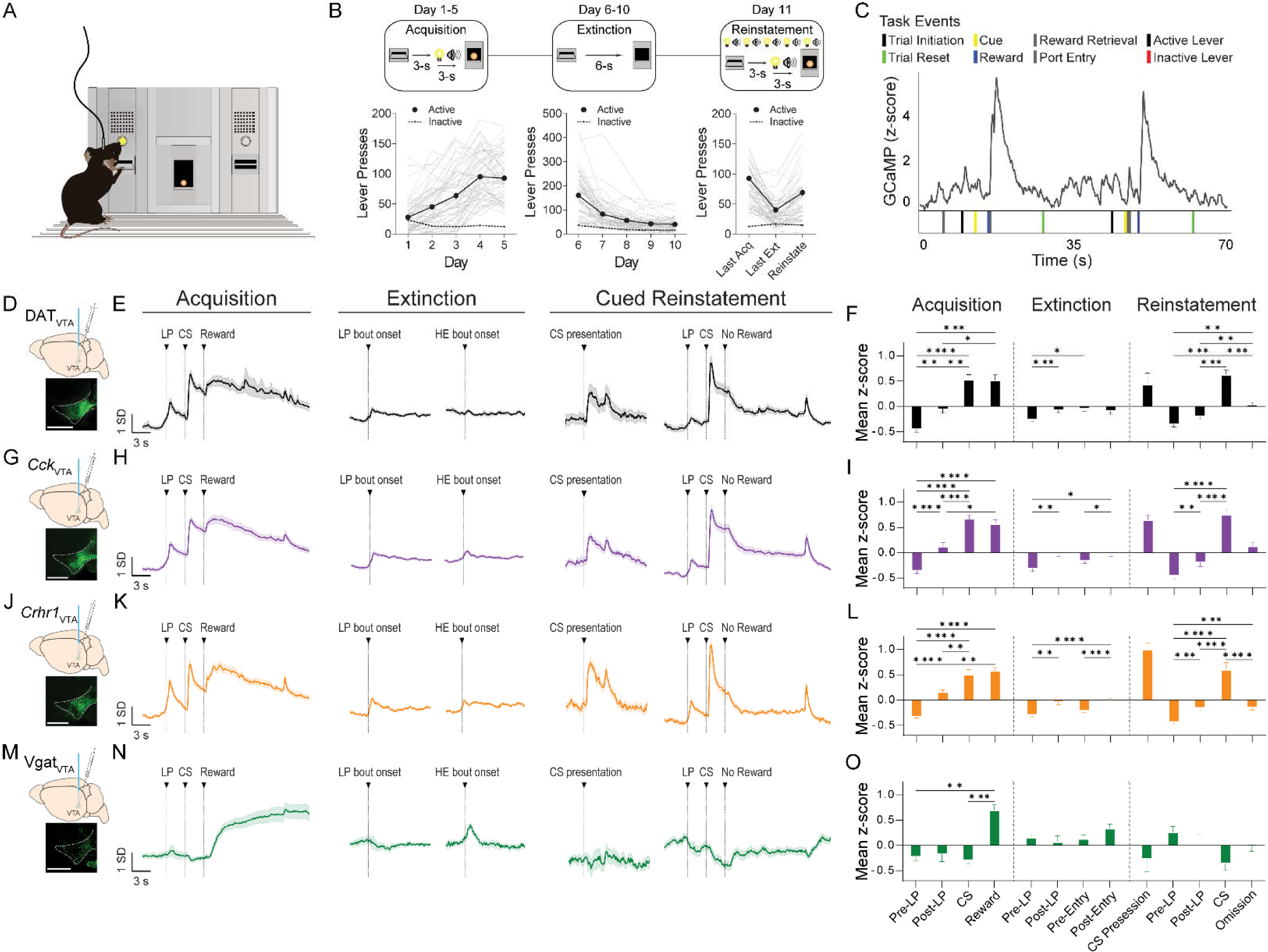
Fiber photometry recordings of VTA subpopulations while mice perform a cued reinstatement task. (A) Schematic of a fiber photometry recording session during the cued reinstatement instrumental conditioning task. (B) Schematic depicting training phases and behavioral performance of mice on acquisition, extinction, and reinstatement task phases (n = 57 mice; solid lines indicate mean across mice and gray lines indicate individual replicates). (C) Example recording trace during acquisition task showing GCaMP fluorescence (top) aligned to event timestamps (bottom). (D) Schematic of viral injection and optic fiber implant and example histology image from the VTA showing staining for GCaMP6 (green) in the DAT_VTA_ group. Scale bar: 500 µm. (E) Z-scored GCaMP fluorescence from DAT_VTA_ population recordings aligned to task events during acquisition, extinction, and reinstatement (n = 9 mice, 63 sessions). Data from all trials during the first, third, and last acquisition and extinction training sessions and from all trials during the reinstatement session. (F) Average z-scored GCaMP fluorescence from DAT_VTA_ population recordings during LP, CS, reward, port entry, and omission periods (n = 9 mice, 63 sessions, bars and error bars indicate mean ± SEM across mice, see Supplementary Table 1 for statistical values). (G) Same as in (D) but for *Cck*_VTA_ population recordings. Scale bar: 500 µm. (H) Same as in (E) but for *Cck*_VTA_ population recordings (n = 16 mice, 112 sessions). (I) Same as in (F) but for *Cck*_VTA_ population recordings (n = 16 mice, 112 sessions). (J) Same as in (D) but for *Crhr1*_VTA_ population recordings. Scale bar: 500 µm. (K) Same as in (E) but for *Crhr1*_VTA_ population recordings (n = 13 mice, 91 sessions). (L) Same as in (F) but for *Crhr1*_VTA_ population recordings (n = 13 mice, 91 sessions). (M) Same as in (D) but for Vgat_VTA_ population recordings. Scale bar: 500 µm. (N) Same as in (E) but for Vgat_VTA_ population recordings (n = 8 mice, 56 sessions). (O) Same as in (F) but for Vgat_VTA_ population recordings (n = 8 mice, 56 sessions).

We next assessed how neural activity in VTA subpopulations correlated with task events (**Figure 1C**). Broadly, the response profile among dopamine populations in response to task-related events was similar but distinct from those observed in Vgat_VTA_ neurons (**Figure 1D-1O**; **Figure S2A-S2E**). GCaMP fluorescence was elevated following reward delivery in all four populations, although the latency to peak following reward delivery was greater in the Vgat_VTA_ population (**Figure 1D-1O**; **Figure S2A**). Interestingly, the Vgat_VTA_ populations also displayed prolonged activity after reward delivery (**Figure 1M-1O**). During acquisition, dopamine populations showed increased activity during the trial-initiation press and CS-presentation periods that was significantly different from the pretrial-initiation (Pre-LP) period (**Figure 1D-1L**). In contrast, GCaMP fluorescence in Vgat_VTA_ neurons during the action and cue periods was not significantly different from the pretrial-initiation period (**Figure 1M-1O**). While both dopamine subpopulations showed phasic responses to these task events, the *Cck*_VTA_ shell-projecting population showed a sustained elevation of GCaMP fluorescence during the full action-cue-outcome period following trial initiation (**Figure 1G-1I**; **Figure S2F**). A comparison between the two dopamine populations revealed a statistically significant difference in the mean response during the action-cue-outcome period across the two populations (**Figure S2F**). In addition, we found *Cck*_VTA_ population activity increased several seconds before mice initiate a trial early in training (**Figure S2G**). By contrast, the *Crhr1*_VTA_ population showed a decrease in activity prior to trial initiation early in training (**Figure S2G**). During extinction, dopamine populations showed modest responses to the lever-press and port-entry bout onsets, while Vgat_VTA_ showed a phasic activation during unrewarded port-entry bout onset (**Figure 1D-1O**). During reinstatement, dopamine populations responded to non-contingent presentations of the cue, but the Vgat_VTA_ population did not (**Figure 1D-1O**). During the contingent phase of reinstatement, dopamine populations showed increased fluorescence to the action and cue responses similar to their response profiles to these periods during acquisition (**Figure 1D-1L**). Vgat_VTA_ neurons showed a sustained decrease in activity during the action and cue periods in contrast to their activity profile during acquisition (**Figure 1M-1O**). During the trial outcome period, the *Cck*_VTA_ population showed a greater latency to decay following omission compared to the *Crhr1*_VTA_ population (**Figure S2H**). Finally, we compared baseline calcium transients during periods of sustained task-related behavioral inactivity during the last extinction session (day 10) across VTA subpopulations (**Figure S1D-S1G**). Consistent with the observed pattern of temporal dynamics during behavioral epochs, the number of transients per minute was significantly greater in the *Crhr1*_VTA_ population (**Figure S1E**) whereas the transient width and amplitude were significantly greater in the *Cck*_VTA_ population (**Figure S1F-S1G**). These results demonstrate that VTA GABA population activity dynamics are largely distinct from those of VTA dopamine subpopulations. Further, dopamine subpopulations show similar neural activity profiles during an instrumental cued reinstatement task, though they display subtle differences in their temporal dynamics during motivated behavior and at baseline.

### Differential encoding of prediction-error and behavioral variables by VTA subpopulations

Our results raise an important question: what do VTA subpopulation responses encode? They may uniformly or preferentially reflect prediction-error and task-related action, cue, and outcome events. Prior work has established that lateral VTA dopamine neurons uniformly encode prediction-errors (Eshel et al., 2016), but how this function is organized across projection-specific VTA subpopulations remains unclear. To test this, we recorded GCaMP fluorescence while mice performed a modified version of the acquisition task in which we introduced random unpredictable reward omissions and reduced the overall reward probability following cue presentation to 50% (**Figure 2A-2B**). Mice showed a significantly shorter latency to initiate a new trial following an unrewarded trial compared to rewarded trials (**Figure 2C**). VTA dopamine populations and the Vgat_VTA_ population showed different responses to reward and omission trials with greater activity following sucrose reward (**Figure 2D-2K**). The VTA dopamine populations show an increase in GCaMP fluorescence following rewarded trial port entries and a decrease in fluorescence following unrewarded port entries (**Figure 2D-2I**). However, latency to the minimum GCaMP response following unrewarded trial port entry was significantly shorter in the *Crhr1*_VTA_ population relative to the *Cck*_VTA_ population (**Figure S3A-S3B**). Additionally, the increase in GCaMP fluorescence following port entry in rewarded trials was greater in the *Cck*_VTA_ neurons relative to the *Crhr1*_VTA_ neurons (**Figure S4C-S4D**). In contrast, the Vgat_VTA_ population shows an increase in fluorescence following both rewarded and unrewarded trial outcomes (**Figure 2J-2K**), consistent with its role in modulating reward prediction in VTA dopamine neurons (Eshel et al., 2015).

**Figure 2.**
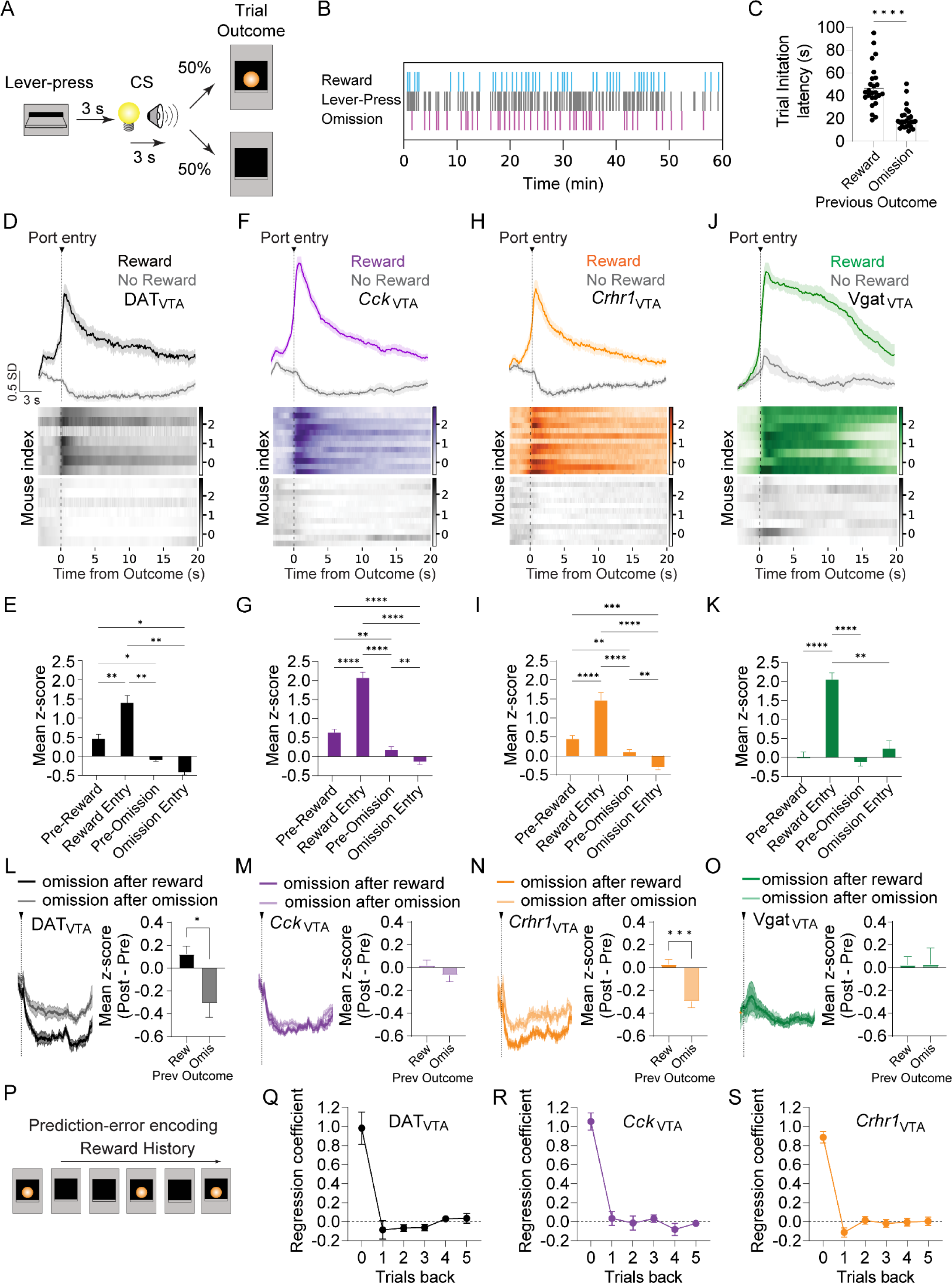
Time-locked activation of VTA subpopulations during random reward omission. (A) Schematic of the random reward-omission task. An active lever-press triggered a 6-s delay-cue period and the probability of reward was 50%. (B) Example behavioral session showing lever-press, reward, and omission times. (C) Trial initiation latency following the intertrial interval period (n = 25 mice, bars and error bars indicate mean ± SEM across mice). (D) Z-scored GCaMP fluorescence and heatmaps aligned to reward (color) or reward omission (gray) task events from the DAT_VTA_ group (n = 7 mice). (E) Average z-scored GCaMP fluorescence during reward, omission, and port entry periods for the DAT_VTA_ (n = 7 mice) group. (F) Same as in (D) but for the *Cck*_VTA_ (n = 12 mice) group. (G) Same as in (E) but for the *Cck*_VTA_ (n = 12 mice) group. (H) Same as in (D) but for the *Crhr1*_VTA_ (n = 13 mice) group. (I) Same as in (E) but for the *Crhr1*_VTA_ (n = 13 mice) group. (J) Same as in (D) but for the Vgat_VTA_ (n = 8 mice) group. (K) Same as in (E) but for the Vgat_VTA_ (n = 8 mice) group. (L) Z-scored GCaMP fluorescence (left) and average z-scored GCaMP fluorescence (right) during reward omission periods shaded according to previous trial outcome type for the DAT_VTA_ group (n = 7 mice). (M) Same as in (L) but for the *Cck*_VTA_ group (n = 12 mice). (N) Same as in (L) but for the *Crhr1*_VTA_ group (n = 13 mice). (O) Same as in (L) but for the Vgat_VTA_ group (n = 8 mice). (P) Schematic of outcome history regression model approach. The current previous five trial outcomes were used with a multiple linear regression to predict the GCaMP signal during the current trial outcome. (Q) Average regression coefficients across mice for the outcome history linear regression for DAT_VTA_ (n = 7 mice), *Cck*_VTA_ neurons (left) (n = 12 mice) and *Crhr1*_VTA_ neurons (right) (n = 13 mice, see Supplementary Table 1 for statistical values). (R) Same as in (Q) but for the *Cck*_VTA_ group (n = 12 mice). (S) Same as in (Q) but for the *Crhr1*_VTA_ group (n = 13 mice).

An essential feature of prediction-error encoding by dopamine neurons is the modulation of reward outcome responses by expectation (Schultz et al., 1997). However, whether prediction-error encoding is uniform across dopamine subpopulations is unknown. First, to compare expectation-dependent modulation of reward-outcome responses among VTA subpopulations, we examined how GCaMP fluorescence correlated with rewarded and unrewarded trial outcomes according to the type of outcome on the preceding trial (**Figure 2L-2O**; **Figure S3E**). In DAT_VTA_ and *Crhr1*_VTA_ populations we observed greater reductions in the GCaMP signal when a reward-omission trial was preceded by a rewarded trial compared to an unrewarded trial (**Figure 2L**; **Figure 2N**). However, we did not observe a change in reward omission response between previous trial outcome conditions in *Cck*_VTA_ neurons (**Figure 2M**). Responses to reward omissions in Vgat_VTA_ neurons were also unaffected by the previous trial outcome (**Figure 2O**). We found that DAT_VTA_ and *Cck*_VTA_ populations showed greater increases in GCaMP fluorescence when a rewarded trial was preceded by an unrewarded trial compared to a rewarded trial (**Figure S2E**). Like *Cck*_VTA_ neurons, we observed significantly greater GCaMP responses in the Vgat_VTA_ population when reward was preceded by and omission (**Figure S2E**). Next, to compare prediction-error encoding in VTA dopamine populations, we fit a linear regression model to predict VTA population trial outcome activity using the current and previous five trial-outcome identities as predictors (**Figure 2P**; adapted from Bayer and Glimcher, 2005). All dopamine populations showed a decay in the influence of previous trial outcomes on their activity (**Figure 2Q-2S**). However, only the DAT_VTA_ and *Crhr1*_VTA_ populations showed the distinctive prediction-error-like combination of a positive modulation by the current trial outcome and a negative modulation by the previous trial outcome (**Figure 2Q**; **Figure 2S**). This result suggests that the *Cck*_VTA_ population preferentially reflects information about current trials whereas the *Crhr1*_VTA_ neurons are modulated by reward history. This is consistent with previous observations that the *Crhr1*_VTA_ population is preferentially involved in reward-related associative learning processes (Heymann et al., 2020).

To test the idea that VTA subpopulations encode distinct information about actions, stimuli, and rewards, we compared the relative contribution of task events (lever-press, cues, outcomes, port entry) to the neural activity in distinct VTA subpopulations. We focused our analysis on the *Cck*_VTA_ and *Crhr1*_VTA_ dopamine subpopulations and the Vgat_VTA_ GABAergic population. To this end, we fit the GCaMP signal of each mouse with a linear encoding model, in which task event type variables are used to predict neural activity (**Figure 3A-3B**; adapted from Engelhard et al., 2019; Parker et al., 2022). To characterize the relative contribution of different task event types in predicting GCaMP fluorescence, we quantified the decrease in explained variance when a given task predictor variable was excluded from the encoding model (**Figure 3C-3H**). Using this approach, we found that the task variable associated with cues showed the greatest relative contribution to the predicted GCaMP signal in the dopamine subpopulations during the action-cue period (**Figure 3C-3D**). Additionally, the *Crhr1*_VTA_ population showed preferential encoding of the cues compared to the trial initiation press (Trial LP) (**Figure 3D**). By contrast, the *Cck*_VTA_ population did not show a significant difference in the relative encoding of cues and the Trial LP (**Figure 3C**). Extraneous lever-press responses that did not initiate a trial reflected in the active (Active LP) and inactive lever-press (Inactive LP) task variables showed smaller relative contributions to predicted neural activity in both dopamine subpopulations (**Figure 3C-3D**). However, the *Cck*_VTA_ population showed preferential encoding of the Trial LP relative to the extraneous active lever-press on the same lever (Active LP) whereas the *Crhr1*_VTA_ population did not show a significant difference in the relative encoding of Trial LP and Active LP task variables (**Figure 3C-3D**). By contrast, none of the task predictor variables had a significantly different relative contribution to neural activity during the action-cue period in the Vgat_VTA_ population (**Figure 3E**). During the trial-outcome period, reward contributed most strongly to the predicted response followed by reward retrieval in both dopamine subpopulations (**Figure 3F-3G**). However, the Vgat_VTA_ population most strongly encoded reward retrieval, followed by reward (**Figure 3H**). Thus, *Cck*_VTA_ and *Crhr1*_VTA_ dopamine subpopulations are distinct in their representation of reward-seeking actions and reward-predictive stimuli but relatively uniform in their encoding of reward outcomes. Further, as a population Vgat_VTA_ neurons do not encode unique information about reward-predictive actions or cue information but strongly encode information about reward receipt. These results indicate that *Cck*_VTA_, *Crhr1*_VTA_, and Vgat_VTA_ dopamine subpopulations encode distinct behavioral variables across action-cue and trial-outcome epochs.

**Figure 3.**
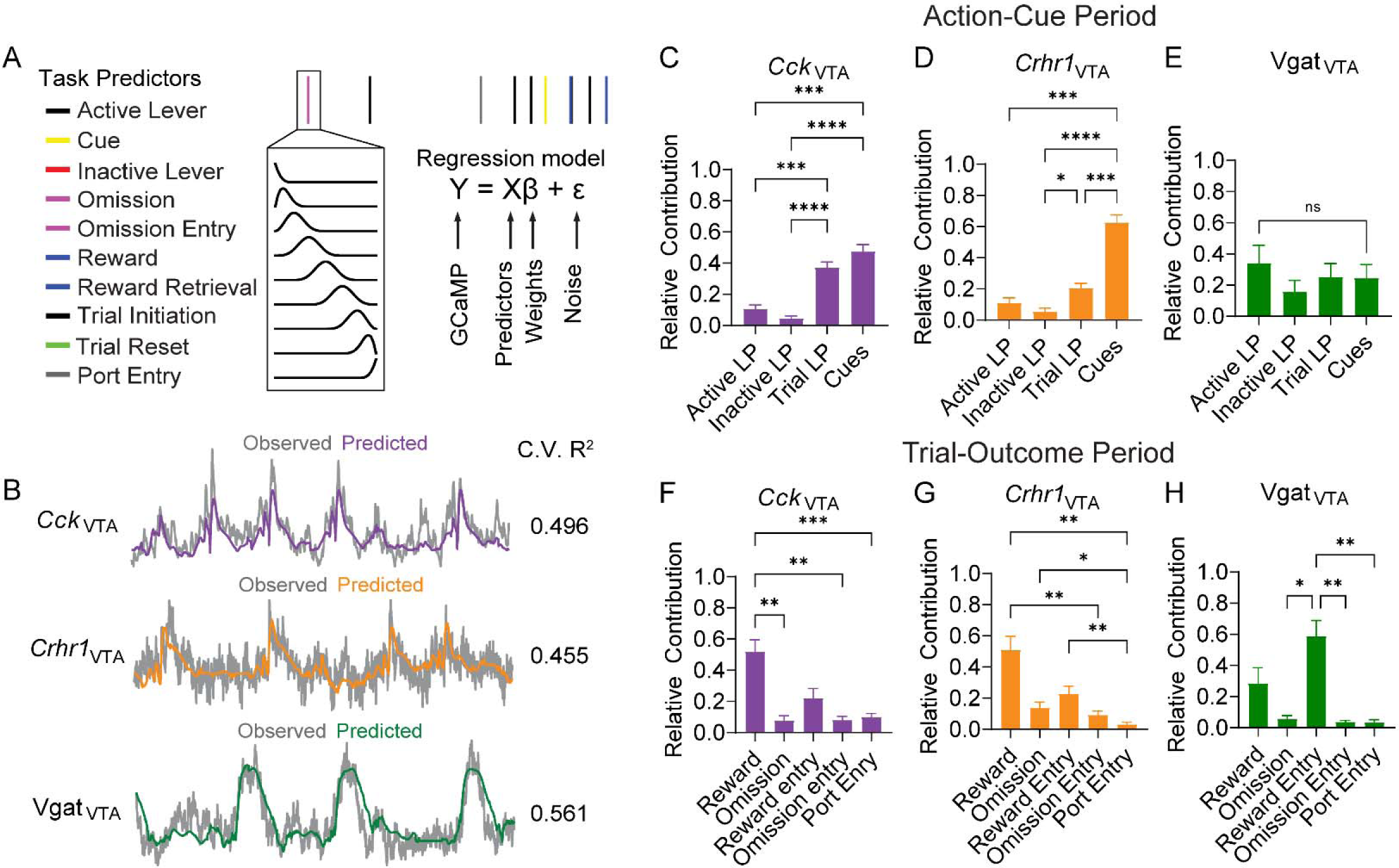
VTA subpopulations differentially encode task relevant behavioral variables during random reward-omission. (A) Schematic of the linear encoding model. Task event timestamps were convolved with a set of cubic splines to generate a predictor set of ten behavior event types. The GCaMP signal was predicted based on task events. (B) Example observed (gray) and predicted (color) GCaMP traces from *Cck*_VTA_ (top), *Crhr1*_VTA_ (middle), and Vgat_VTA_ (bottom) groups. (C) Relative contribution of each task event type to the explained variance of the GCaMP signal during the action-cue period, averaged across mice for the *Cck*_VTA_ group (n = 12 mice) (see Supplementary Table 1 for statistical values). (D) Same as in (C) but for the *Crhr1*_VTA_ group (n = 11 mice). (E) Same as in (C) but for the Vgat_VTA_ group (n = 8 mice). (F) Relative contribution of each task event type to the explained variance of the GCaMP signal during the trial outcome period, averaged across mice for the *Cck*_VTA_ group (n = 12 mice). (G) Same as in (F) but for the *Crhr1*_VTA_ group (n = 11 mice). (H) Same as in (F) but for the Vgat_VTA_ group (n = 8 mice).

### VTA populations differentially encode reward anticipation

The above results demonstrate how VTA subpopulations respond when rewards are readily attainable and the response requirement is fixed. However, VTA dopamine neuron activity is correlated with effort exertion and dopamine release in the NAc is thought to invigorate ongoing motivated behaviors (Salamone and Correa, 2012; Gan et al., 2010). How dynamic changes in effort and reward availability are represented across VTA dopamine subpopulations and the VTA GABA population remains unknown. To address this question, we recorded GCaMP fluorescence while mice underwent a progressive ratio test, in which the number of lever-presses required to earn a reward is increased systematically following each reward throughout the session (**Figure 4A**). This task also measures the breakpoint, or the maximum level of effort an animal is willing to exert before they stop responding (Hodos, 1961). Mice tracked this increasing response requirement, reflected in the high number of cumulative lever-press responses across mice (**Figure 4B-4C**). The increasing response requirement allowed us to measure responses reflective of relative levels of effort exertion and reward anticipation across decreasing reward availability periods.

**Figure 4.**
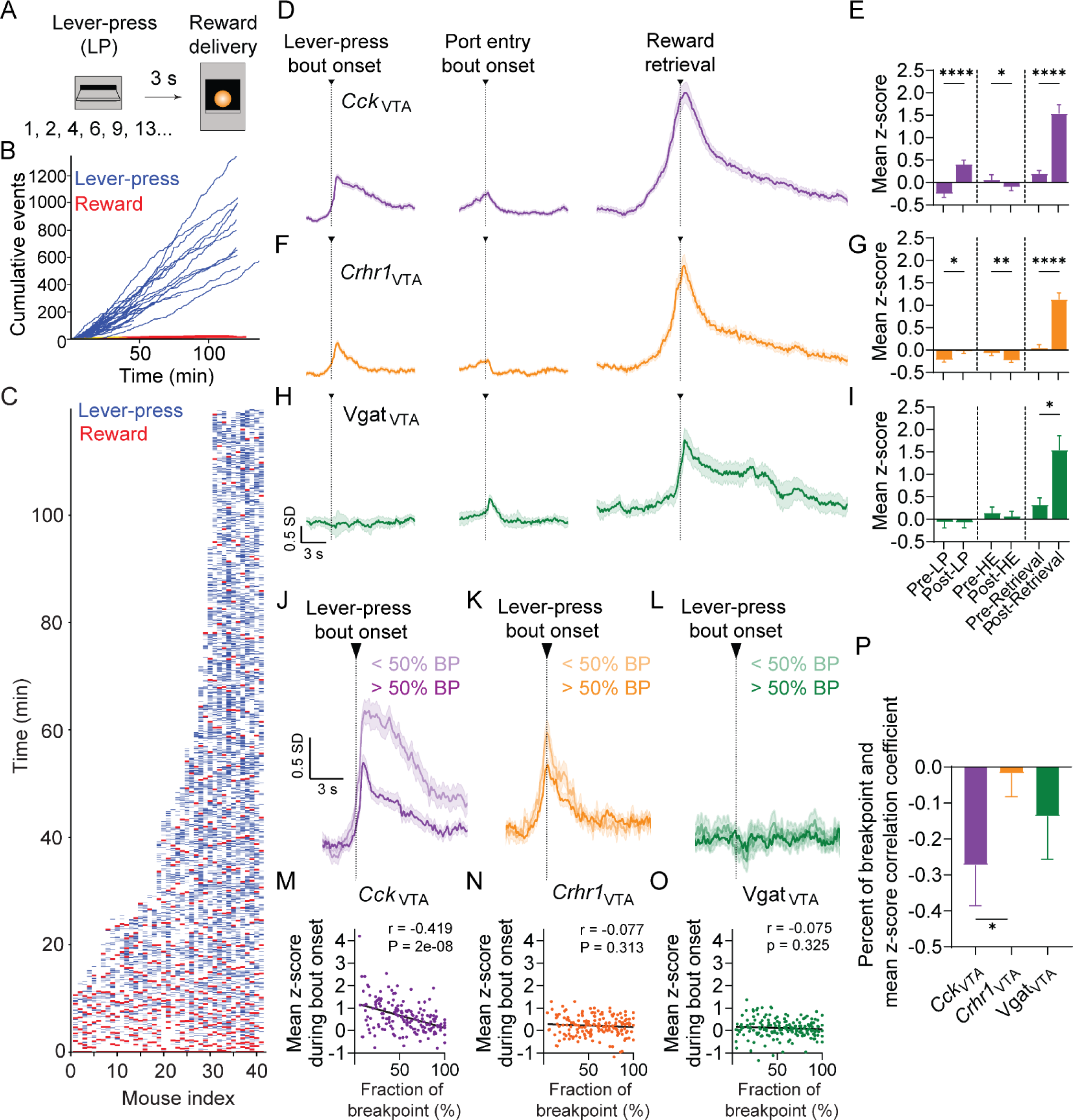
VTA subpopulations differentially encode reward anticipation during progressive ratio. (A) Schematic of the progressive ratio (PR) task. The number of lever-presses required for a reward increased systematically following each reward throughout the session. (B) Cumulative lever-presses (blue) and rewards (red) during PR sessions. Lines indicate individual mice (n = 42 mice). (C) Lever-press (blue) and reward (red) event times shown for all sessions (n = 42 mice). (D) Z-scored GCaMP fluorescence from photometry recordings aligned to lever-press bout onset, port entry bout onset, and reward retrieval for *Cck*_VTA_ group (n = 17 mice). (E) Average z-scored GCaMP fluorescence during lever-press bout onset, port entry bout onset, and reward retrieval periods for the *Cck*_VTA_ group (n = 17 mice). Bars and error bars indicate mean ± SEM across mice. (F) Same as in (D) but for the *Crhr1*_VTA_ group (n = 17 mice). (G) Same as in (E) but for the *Crhr1*_VTA_ group (n = 17 mice). (H) Same as in (D) but for the Vgat_VTA_ group (n = 8 mice). (I) Same as in (E) but for the Vgat_VTA_ group (n = 8 mice). (J) Z-scored GCaMP fluorescence from recordings aligned to lever-press bout onset, separated by bouts occurring prior to (lighter shade) and after (darker shade) 50% of all completed reinforcement ratios for the *Cck*_VTA_ group (n = 17 mice). (K) Same as in (J) but for the *Crhr1*_VTA_ group (n = 17 mice). (L) Same as in (J) but for the Vgat_VTA_ group (n = 8 mice). (M) Correlations across bouts between percent of breakpoint and mean z-scored GCaMP signal during bout onset for the *Cck*_VTA_ group (n = 165 bouts). Correlation coefficient (r) and p-values on the top right of the plot. (N) Same as in (M) but for the *Crhr1*_VTA_ group (n = 172 bouts). (O) Same as in (M) but for the Vgat_VTA_ group (n = 172 bouts). (P) Pearson’s correlation coefficient per mouse between percent breakpoint and mean z-scored GCaMP signal for *Cck*_VTA_ (n = 17 mice), *Crhr1*_VTA_ (n = 17 mice), and Vgat_VTA_ (n = 8 mice) groups (see Supplementary Table 1 for statistical values).

Across the entire session, we found that activity in both dopamine subpopulations is transiently increased at the onset of bouts of lever-pressing (**Figure 4D-4G**). In addition, the *Cck*_VTA_ population showed a sustained elevation in activity during the 10-s following bout onset (**Figure 4D-4E**). By contrast, the Vgat_VTA_ population was not engaged at lever-press bout onset (**Figure 4H-4I**). During unrewarded port entry bout onset all VTA populations showed a small increase in activity prior to bout onset followed by a decrease to baseline during the duration of the bout (**Figure 4D-4I**). During rewarded port entries activity in both dopamine subpopulations ramped up prior to reward retrieval and shows sustained activity during reward consumption (**Figure 4D-4G**). Vgat_VTA_ neurons showed a relatively modest increase in activity prior to reward retrieval, in contrast to VTA dopamine populations (**Figure 4H-4I**).

To determine how the response profiles of VTA subpopulations evolve across increasing effort requirements and decreasing reward availability, we examined neural activity as a function of the breakpoint of each mouse. We found that *Crhr1*_VTA_ and *Cck*_VTA_ populations showed distinct activity profiles in response to decreasing reward availability. *Cck*_VTA_ population activity at lever-press bout onset was initially high and sustained during the entire bout earlier in the session with lower lever-press response requirements (**Figure 4J**). However, *Cck*_VTA_ population activity during lever-press bout onset was decreased later in the session as rewards required more effort exertion to obtain them (**Figure 4J**). By contrast, the *Crhr1*_VTA_ population responded similarly at lever-press bout onset regardless of response requirements (**Figure 4K**). Thus, the *Cck*_VTA_ population is preferentially engaged with sustained activity during ongoing motivated responding when rewards are more readily attainable. Further, we found that decreased reward availability was associated with less activity in the *Cck*_VTA_ population during lever-press bouts across the entire session but not in the *Crhr1*_VTA_ or Vgat_VTA_ population (**Figure 4M-4O**). A statistical comparison of the correlation between percent of breakpoint and GCaMP response during LP bout onset across mice revealed a significant difference between the *Cck*_VTA_ and *Crhr1*_VTA_ populations (**Figure 4P**). Taken together, these results suggest that *Cck*_VTA_ dopamine population activity reflects a reward anticipation signal during goal-directed actions that tracks reward availability, whereas the *Crhr1*_VTA_ dopamine population does not encode this type of motivational information. This is consistent with the previous observation that the *Cck*_VTA_ population is preferentially involved in reward-driven motivated responding.

### Differential contribution of dopamine subpopulations to cued reinstatement of reward-seeking behavior

*Crhr1*_VTA_ and *Cck*_VTA_ dopamine subpopulations differentially regulate the acquisition and extinction of an instrumental response (Heymann et al., 2020). Further, it has been shown that phasic activation of VTA dopamine neurons is sufficient to reactivate a previously extinguished instrumental behavior in the absence of reward-predictive stimuli (Tsai et al., 2009; Adamantidis et al., 2011; Witten et al., 2011). However, how distinct VTA dopamine subpopulations mediate the reactivation of cue-driven instrumental responding following extinction remains unclear. To determine if reward-predictive, cue-related neural activity in dopamine subpopulations is sufficient to modulate the reactivation of cue-driven instrumental responding, we photostimulated both populations during CS-presentation throughout the reinstatement session. *Cck*-Cre, and *Crhr1*-Cre mice were injected with an AAV expressing a Cre-dependent ChR2 or GFP in the VTA and implanted with an optical fiber above the VTA (**Figure 5A**; **Figure S4A**). Mice were trained on the acquisition and extinction phases of the cued reinstatement task without photostimulation (**Figure 5B-5C**; **Figure S4B-S4C**). Extinction-resistant mice were excluded from the analysis which resulted in the exclusion of 1 of 17 controls, 1 of 22 *Crhr1-*Cre, and 2 of 18 *Cck-Cre* mice. To rule out the possible confound of reinforcement of lever-pressing by photostimulation in the absence of CS-presentation, we photostimulated each population 3-s (20 Hz, 5-ms pulse width, 3-s duration) following an active lever-press during an additional extinction session (Stim) prior to the reinstatement session (Stim + CS) (**Figure 5B**). Photostimulation of either population 3-s following an active lever-press in the absence of CS-presentation did not affect operant responding (**Figure 5D**). We next asked whether photostimulation of these neurons during CS-presentation is sufficient to alter reinstatement behavior (Stim + CS) (**Figure 5B**). CS-presentation paired with photostimulation of *Crhr1*_VTA_ but not *Cck*_VTA_ neurons during cued reinstatement significantly increased the number of active lever-presses during the entire session (**Figure 5D-5E**) but did not alter the total number of trials completed (**Figure 5F**) or the number of inactive lever-presses (**Figure S4D**).

**Figure 5.**
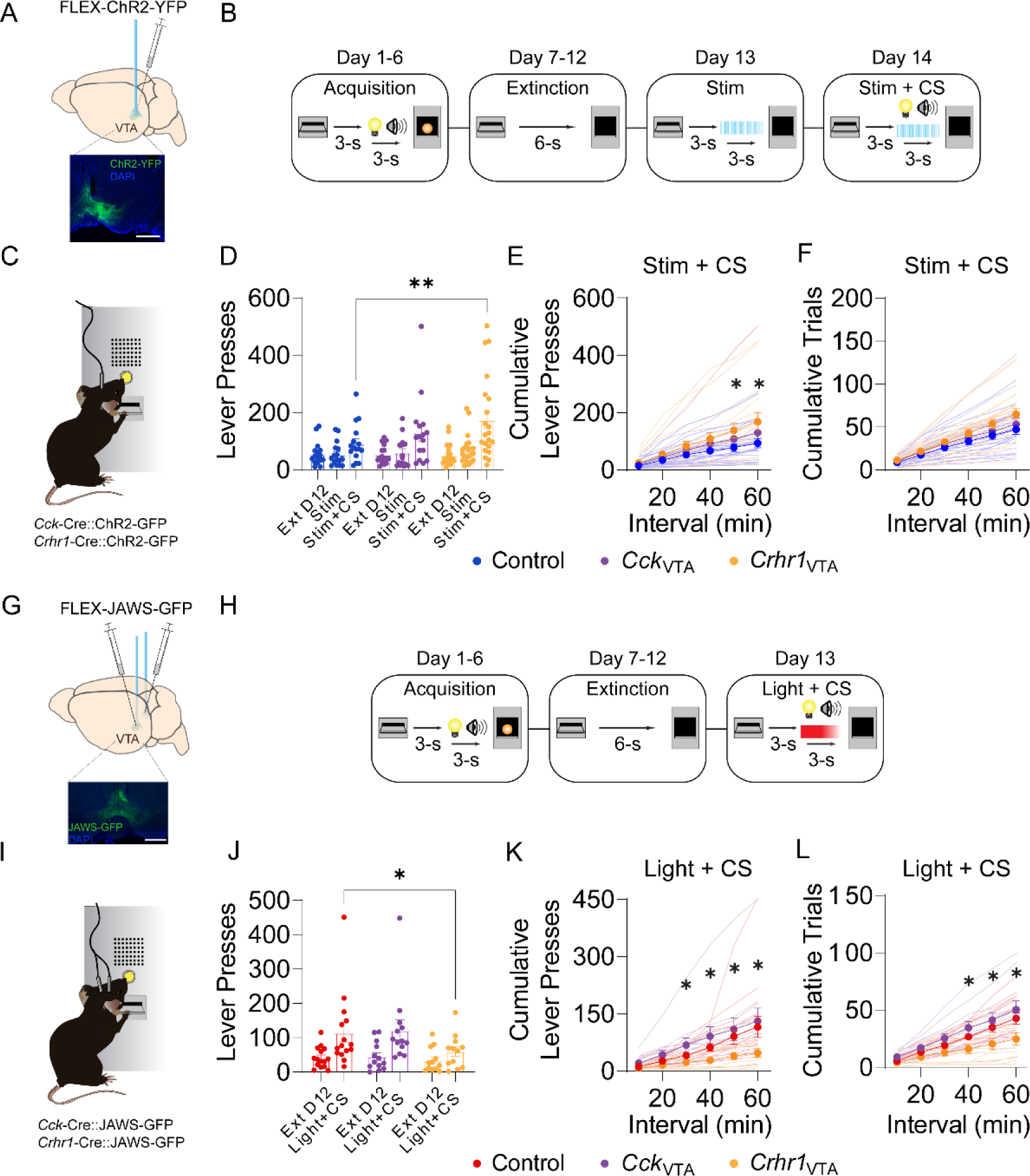
Photostimulation and photoinhibition of *Crhr1*_VTA_ and *Cck*_VTA_ neurons during cued reinstatement. (A) Schematic of an optogenetic cued reinstatement session with channelrhodopsin (ChR2) stimulation. (B) Schematic of the training phases of the optogenetic cued reinstatement task. On day 13 an active lever-press triggered a 3 s delay followed by blue light stimulation. On day 14 an active lever-press triggered a 3 s delay followed by blue light stimulation paired with CS presentation. (C) Schematic of viral injection and optic fiber implant and example histology image from the VTA showing staining for ChR2-YFP (green) in VTA subpopulations. (D) Mean number of lever-presses across mice during acquisition and extinction sessions in control, *Cck*_VTA_, and *Crhr1*_VTA_ groups (n = 11-21 mice, error bars represent SEM). (E) Mean number of cumulative lever-presses on day 14 (Stim + CS) with optogenetic activation, or in control mice without opsin expression (n = 11-21 mice, error bars indicate SEM). (F) Mean number of cumulative trials completed on day 14 (Stim + CS) with optogenetic activation, or in control mice without opsin expression (n = 11-21 mice, error bars indicate SEM). (G) Schematic of an optogenetic cued reinstatement session with JAWS-GFP (JAWS) stimulation. (H) Schematic of the training phases of the optogenetic cued reinstatement task. On day 13 an active lever-press triggered a 3 s delay followed by red light stimulation paired with CS presentation. (I) Schematic of viral injection and optic fiber implant and example histology image from the VTA showing staining for JAWS-GFP (green) in VTA subpopulations. (J) Mean number of lever-presses on day 12 (Ext D12), and day 13 (Light + CS) with optogenetic inhibition, or in control mice without opsin expression (n = 12-14 mice, error bars indicate SEM). (K) Mean number of cumulative lever-presses on day 13 (Light + CS) with optogenetic inhibition, or in control mice without opsin expression (n = 12-14 mice, error bars indicate SEM). (L) Mean number of cumulative trials completed on day 13 (Light + CS) with optogenetic inhibition, or in control mice without opsin expression (n = 12-14 mice, error bars indicate SEM, see Supplementary Table 1 for statistical values).

To determine if the activity observed in dopamine subpopulations during CS-presentation causally contributes to cued reinstatement behavior, we photoinhibited both populations during CS-presentation throughout the reinstatement session (**Figure 5H**). *Cck-*Cre and *Crhr1-*Cre mice were injected bilaterally with an AAV carrying Cre-dependent inhibitory JAWS or GFP in the VTA and implanted with bilateral optical fibers above the VTA (**Figure 5G**; **Figure S4E**). Mice performed the acquisition and extinction phases of the cued reinstatement task without photoinhibition (**Figure 5H-5I**; **Figure S4F-S4G**). No mice met exclusion criteria for extinction resistance. VTA subpopulations were photoinhibited 3-s (2-s constant square pulse terminated with a 1-s linear ramp-down to avoid rebound excitation, Jo et al., 2018) during the CS-presentation throughout the cued reinstatement session (Light + CS) (**Figure 5H**). Photoinhibition of *Crhr1*_VTA_ but not *Cck*_VTA_ neurons reduced the total number of lever-presses in the reinstatement session (Light + CS) (**Figure 5J-5K**) and reduced the total number of trials completed (**Figure 5L**). Photoinhibition did not alter the number of inactive lever-presses (**Figure S4H**). Taken together, these findings indicate that the contribution of *Crhr1*_VTA_ neurons to cued reinstatement is distinct from that of *Cck*_VTA_ neurons.

### Baseline electrophysiology of Crhr1_VTA_ and Cck_VTA_ populations

We also examined whether intrinsic electrophysiological properties could underlie the differential response profiles of *Cck*_VTA_ and *Crhr1*_VTA_ neurons during behavior. To label neurons for recording, we injected *Cck*-Cre and *Crhr1*-Cre mice with an AAV carrying Cre-dependent eYFP in the VTA (**Figure 6A**). We performed whole-cell, voltage and current-clamp recording *ex vivo* and found distinct properties of excitability and inhibitory transmission in *Cck*_VTA_ and *Crhr1*_VTA_ neurons (**Figure 6B-6J**). To examine the intrinsic excitability of Crhr1_VTA_ and Cck_VTA_ subpopulations, we recorded action potential firing from eYFP-labeled cells in response to increasing steps of current injection (**Figure 6B-6D**). *Cck*_VTA_ and *Crhr1*_VTA_ neurons continued to increase firing with increasing current injections (**Figure 6C**). However, *Cck*_VTA_ neurons fired a greater number of action potentials in response to depolarizing steps of current and showed a shorter latency to spike following current injection compared to *Crhr1*_VTA_ neurons (**Figure 6C-6D**). These results revealed an increased excitability of *Cck*_VTA_ neurons which could contribute to their sustained activity profiles during behavior.

**Figure 6.**
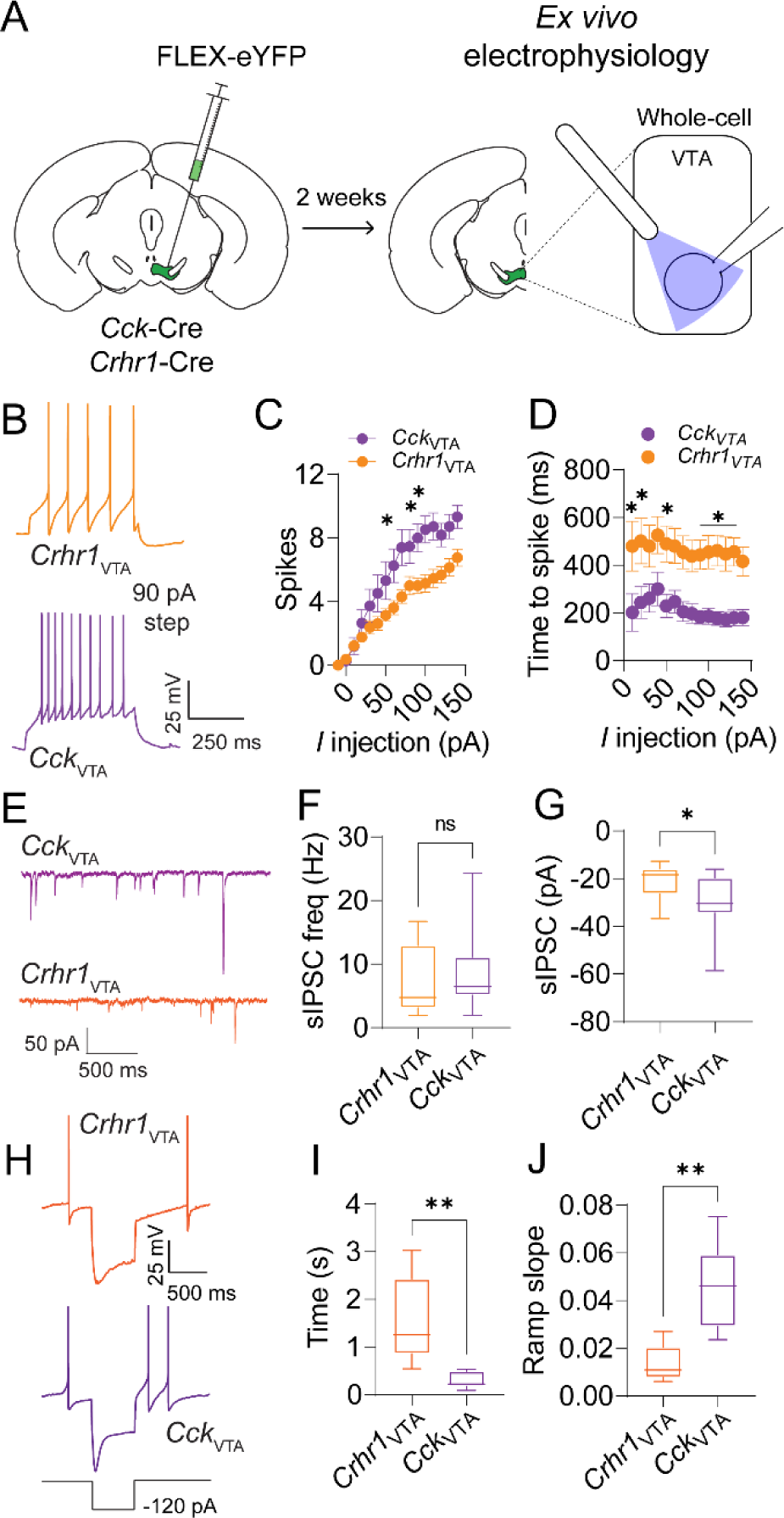
Baseline neurophysiological properties *Crhr1*_VTA_ and *Cck*_VTA_ neurons. (A) Schematic of the viral injection strategy and *ex vivo* electrophysiology recordings. (B) Representative evoked excitability traces from *Crhr1*_VTA_ (orange) or *Cck*_VTA_ (magenta) neurons (90 pA current injection). (C) Current-voltage plot for *Crhr1*_VTA_ (orange) and *Cck*_VTA_ (magenta) neurons (n = 13-15 cells) showing mean number of evoked spikes per current injection level. Bars and error bars indicate mean ± SEM across cells. (D) Mean spike latency following current injection for *Crhr1*_VTA_ (orange) and *Cck*_VTA_ (magenta) neurons (n = 13-15). Bars and error bars indicate mean ± SEM across cells. Representative spontaneous inhibitory postsynaptic current (sIPSC) traces from *Crhr1*_VTA_ (orange) or *Cck*_VTA_ (magenta) neurons. (E) Mean sIPSC frequency for *Crhr1*_VTA_ (orange) and *Cck*_VTA_ (magenta) neurons (n = 13-15 cells). (F) Mean sIPSC amplitude for *Crhr1*_VTA_ (orange) and *Cck*_VTA_ (magenta) neurons (n = 13-15 cells). (G) Representative traces of rebound spiking from *Crhr1*_VTA_ (orange) or *Cck*_VTA_ (magenta) neurons following injection of a -120 pA hyperpolarizing current. (H) Mean time to first spike following hyperpolarization for *Crhr1*_VTA_ (orange) and *Cck*_VTA_ (magenta) neurons (n = 5-6 cell) (I) Mean ramp slope prior to first spike following hyperpolarization for *Crhr1*_VTA_ (orange) and *Cck*_VTA_ (magenta) neurons (n = 5-6 cells, see Supplementary Table 1 for statistical values).

We then performed voltage-clamp recordings and recorded spontaneous inhibitory postsynaptic currents (sIPSCs), which reflect spontaneous neurotransmitter release (**Figure 6E**). *Cck*_VTA_ and *Crhr1*_VTA_ neurons did not differ in sIPSC frequency (**Figure 6F**), suggesting that these subpopulations share similar mechanisms of presynaptic inhibitory transmission. However, compared with *Crhr1*_VTA_ neurons, *Cck*_VTA_ neurons showed greater sIPSC amplitude (**Figure 6G**), indicating that these subpopulations differ in their postsynaptic mechanisms of inhibitory synaptic transmission. Finally, we asked if there is a difference in rebound spiking following injection of a hyperpolarizing current between *Cck*_VTA_ and *Crhr1*_VTA_ neurons. *Cck*_VTA_ neurons showed a shorter latency to spike and a faster membrane potential rise time following the offset of a hyperpolarizing step compared to *Crhr1*_VTA_ neurons (**Figure 6H-6J**). Taken together, these results establish distinct intrinsic membrane properties of *Cck*_VTA_ and *Crhr1*_VTA_ neurons, which could contribute to their distinct activity patterns *in vivo*.

### Functional optogenetic characterization of inhibitory and disinhibitory inputs to *Cck*_VTA_ and *Crhr1*_VTA_ populations

Diverse inhibitory and disinhibitory connections play important roles in the regulation of VTA dopamine neuron activity and motivated behavior (Morales and Margolis, 2017). Previous work has established that VTA GABA neurons directly control VTA dopamine neuron excitability and ongoing motivated behavior (Van Zessen et al., 2012). Additionally, two prominent GABAergic inputs to the VTA from the lateral hypothalamus (LH) and nucleus accumbens medial shell (NAc mshell) form disinhibitory connections with VTA dopamine neurons via the VTA GABA population and differentially regulate dopamine neuron activity, immediate early gene activation, and motivated behavior (Nieh et al., 2016; Yang et al., 2018; Soden et al., 2020; Simon et al., 2023). However, how VTA GABAergic neurons and GABAergic inputs to the VTA regulate the activity of distinct dopamine subpopulations remains unresolved.

We hypothesized that activation of VTA GABA neurons, LH GABA neurons, or NAc mshell GABA neurons would drive distinct response profiles in *Cck*_VTA_ and *Crhr1*_VTA_ dopamine neurons. To test this possibility, we photostimulated either Vgat_VTA_ neurons, LH GABA neurons, or NAc mshell GABA neurons while recording the neural activity of VTA dopamine subpopulations. First, we injected *Cck*-Cre::*Vgat*-Flp or *Crhr1*-Cre::*Vgat*-Flp mice with an AAV carrying Flp-dependent ChrimsonR-tdTomato in Vgat_VTA_ neurons and an AAV carrying Cre-dependent GCaMP6m in either *Cck*_VTA_ and *Crhr1*_VTA_ neurons and implanted an optic fiber for dual recording and stimulation above the VTA (**Figure 7A**; **Figure S5A**). In freely moving mice, we photostimulated the Vgat_VTA_ population with red light (20 Hz or 40 Hz, 3-s duration) and observed a significant decrease in GCaMP fluorescence relative to the pre-stimulation period in both *Cck*_VTA_ and *Crhr1*_VTA_ populations (**Figure 7B**). However, compared to the *Cck*_VTA_ population, the *Crhr1*_VTA_ population showed a larger amplitude inhibitory response (**Figure 7B**). This suggests that local inhibition may control the activity of *Crhr1*_VTA_ neurons more strongly *in vivo*.

**Figure 7.**
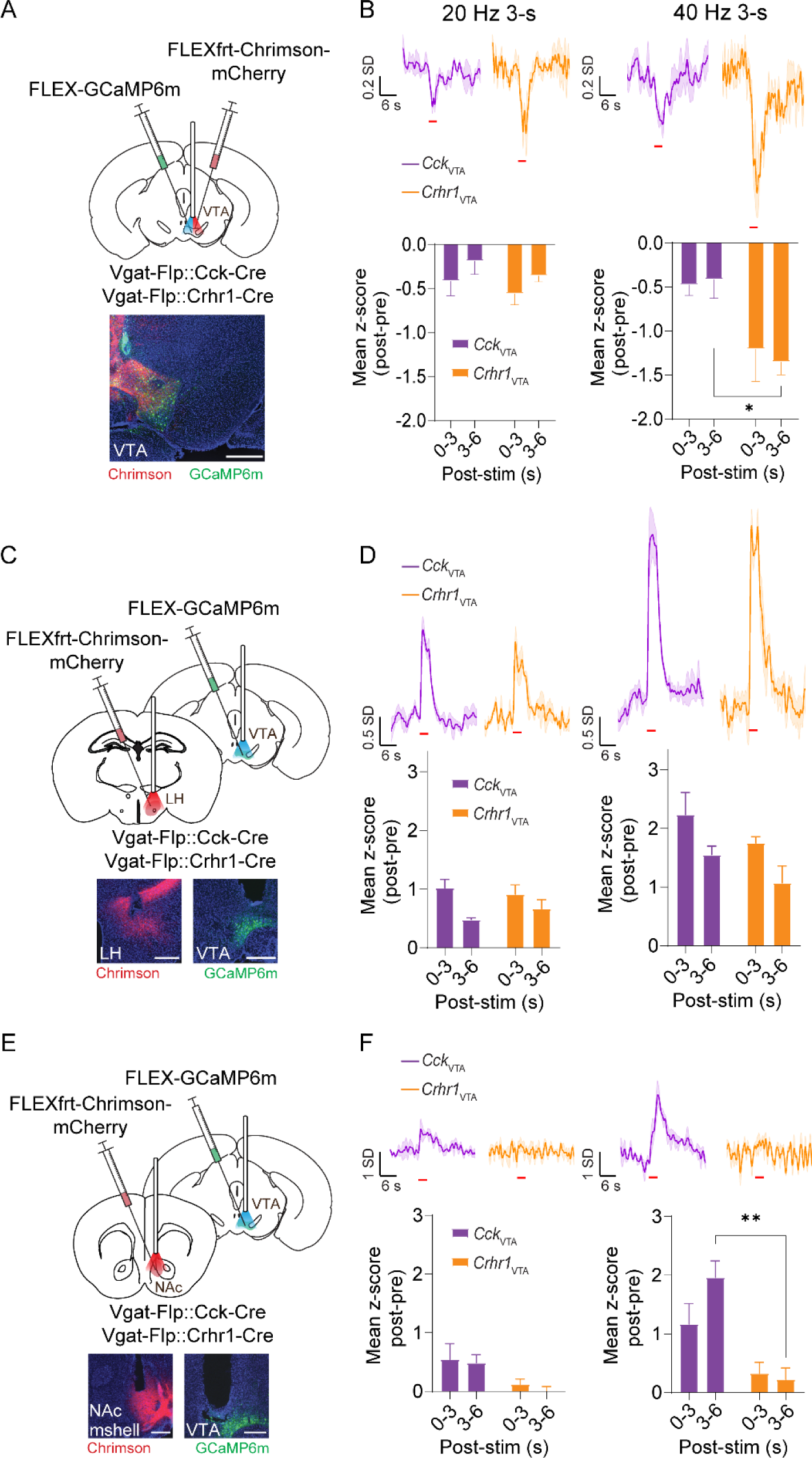
Functional optogenetic characterization of inhibitory and disinhibitory inputs to *Crhr1*_VTA_ and *Cck*_VTA_ populations. (A) Schematic of viral injection strategy. Vgat-Flp::*Cck*-Cre or Vgat-Flp::*Crhr1*-Cre mice were injected with Flp-dependent Chrimson and Cre-dependent GCaMP into the VTA. An optical fiber for dual stimulation and recording was implanted above VTA. Scale bar: 500 µm. (B) Z-scored GCaMP fluorescence aligned to Vgat_VTA_ stimulation in *Cck*_VTA_ (n = 3 mice, 6 sessions) and *Crhr1*_VTA_ (n = 3 mice, 6 sessions) groups (top). Average z-scored GCaMP fluorescence during stimulation and post-stimulation periods for *Cck*_VTA_ (n = 3 mice, 10 sessions) and *Crhr1*_VTA_ (n = 3 mice, 6 sessions) groups (bottom). Bars and error bars indicate mean ± SEM across mice. (C) Schematic of viral injection strategy. Vgat-Flp::*Cck*-Cre or Vgat-Flp::*Crhr1*-Cre mice were injected with Flp-dependent Chrimson into the LH and Cre-dependent GCaMP into the VTA. Stimulation and recording fibers were implanted above the LH and VTA, respectively. Scale bar: 500 µm. (D) Z-scored GCaMP fluorescence aligned to LH GABA stimulation in *Cck*_VTA_ (n = 5 mice, 10 sessions) and *Crhr1*_VTA_ (n = 3 mice, 6 sessions) groups (top). Average z-scored GCaMP fluorescence during stimulation and post-stimulation periods for *Cck*_VTA_ (n = 5 mice, 10 sessions) and *Crhr1*_VTA_ (n = 3 mice, 6 sessions) groups (bottom). Bars and error bars indicate mean ± SEM across mice. (E) Schematic of viral injection strategy. Vgat-Flp::*Cck*-Cre or Vgat-Flp::*Crhr1*-Cre mice were injected with Flp-dependent Chrimson into the NAc mshell and Cre-dependent GCaMP into the VTA. Stimulation and recording fibers were implanted above the NAc mshell and VTA, respectively. Scale bar: 500 µm. (F) Z-scored GCaMP fluorescence aligned to NAc shell stimulation in *Cck*_VTA_ (n = 3 mice, 6 sessions) and *Crhr1*_VTA_ (n = 3 mice, 6 sessions) groups (top). Average z-scored GCaMP fluorescence during stimulation and post-stimulation periods for *Cck*_VTA_ (n = 3 mice, 6 sessions) and *Crhr1*_VTA_ (n = 3 mice, 6 sessions) groups (bottom). Bars and error bars indicate mean ± SEM across mice (see Supplementary Table 1 for statistical values).

We next examined how stimulation of the LH GABA or NAc mshell GABA population drives activity in VTA dopamine subpopulations. Using a similar strategy, we expressed Flp-dependent ChrimsonR-tdTomato in either LH GABA neurons or NAc mshell GABA neurons and GCaMP6m in either *Cck*_VTA_ or *Crhr1*_VTA_ neurons and implanted an optic fiber for stimulation above the LH or NAc mshell and an optic fiber for recording above the VTA (**Figure 7C**; **Figure 7E**; **Figure S5B-S5E**). We found that photostimulation of LH GABA strongly activated both VTA dopamine populations (**Figure 7D**). In contrast, photostimulation of NAc mshell GABA evoked a sustained activation only in the *Cck*_VTA_ population at both stimulation frequencies (**Figure 7F**). Thus, *Cck*_VTA_ and *Crhr1*_VTA_ populations respond differentially to local VTA GABAergic and disinhibitory VTA GABAergic inputs. Taken together, these findings support a model in which differential inhibitory connectivity among dopamine subpopulations contributes to heterogeneous response profiles *in vivo*.

### Mapping brain-wide monosynaptic inputs to VTA dopamine subpopulations

VTA dopamine neurons receive synaptic input from numerous brain regions (Watabe-Uchida et al., 2012; Ogawa et al., 2014; Faget et al., 2016; Chung et al., 2017) and distinct VTA dopamine projection populations are differentially regulated by multiple upstream regions (Lammel et al., 2012, Beier et al., 2015). We hypothesized that *Cck*_VTA_ neurons and *Crhr1*_VTA_ neurons receive distinct patterns of monosynaptic input across the whole brain. To test this, we compared the relative density of brain-wide monosynaptic inputs to *Cck*_VTA_ and *Crhr1*_VTA_ populations using rabies virus-based transsynaptic retrograde tracing (Sun et al., 2014) paired with tissue clearing and light sheet fluorescent microscopy (LSFM). AAV-syn-DIO-TC66T-2A-eGFP-2A-oG was injected into the VTA of *Cck*-Cre or *Crhr1*-Cre mice followed by injection of EnvA-SADΔG-RV-DsRed 14 days later (**Figure 8A**). After 9 days, intact brains were optically cleared and imaged using LSFM (**Figure 8A**). Starter cell populations in the VTA were identified based on eGFP and DsRed coexpression (**Figure S6A**). Transsynaptically labeled neurons were identified based on DsRed-only expression (**Figure S6B**). To quantify the anatomical distribution of input cells, we used a modified ClearMap pipeline (Renier et al., 2016; Madangopal et al., 2022) for brain atlas registration and automated cell detection. For both *Cck*_VTA_ and *Crhr1*_VTA_ starter cell populations, DsRed-positive input cells were found across the brain (**Figure 8B-8D**; **Figure S6B-6D**; **Figure S7**). While the total number input cells varied across mice, the number of starter cells and input cells was roughly proportional (**Figure S6E**). Cell counts across all brain regions were normalized to the total number of input cells for each mouse to account for variability in the total number of labeled neurons.

**Figure 8.**
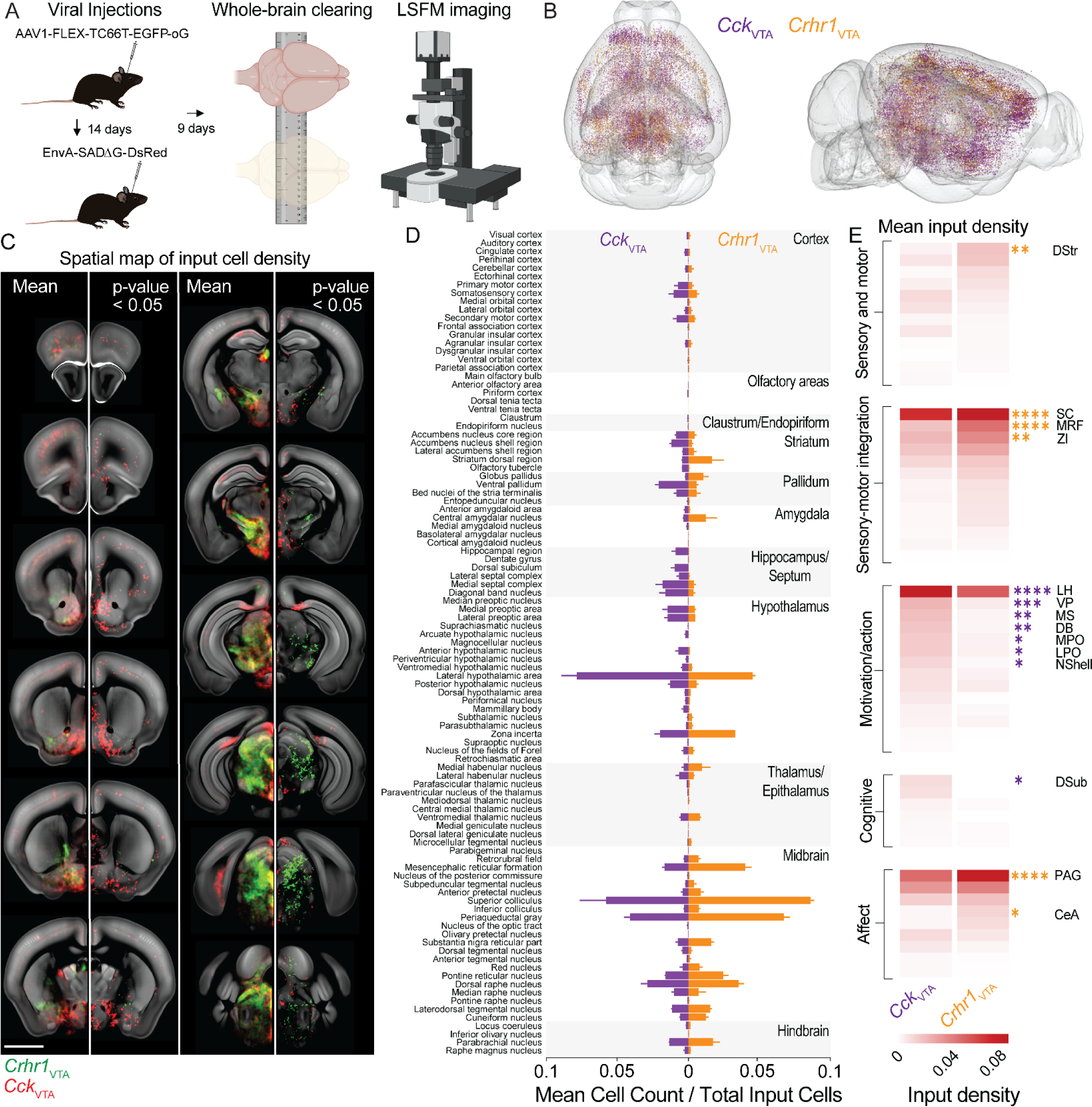
Whole-brain mapping of inputs to *Crhr1*_VTA_ and *Cck*_VTA_ neurons. (A) Schematic of viral injection strategy, whole-brain clearing, and light sheet fluorescence microscopy (LSFM). Cre-dependent helper virus (AAV-syn-DIO-TC66T-2A-eGFP-2A-oG) was injected into the VTA of *Cck*-Cre or *Crhr1*-Cre mice. Two weeks later, rabies virus (EnvA-SADΔG-RV-DsRed) was injected in to the VTA. Nine days later, intact brains were cleared and imaged. (B) Location of input cells to *Cck*_VTA_ and *Crhr1*_VTA_ neurons in example mice. (C) Voxelized heatmap of input cell density in coronal sections across the whole-brain. Mean density of DsRed-positive cells per mm^3^ across mice (left panels). Voxelized results of group one-way ANOVA pairwise comparison (right panels) (n = 3 mice). Scale bar: 2 mm. (D) Mean number of input cells normalized to total number of input cells for all input regions for *Cck*_VTA_ (left) and *Crhr1*_VTA_ (right) groups (n = 3 mice). (E) Heatmap of group pairwise comparison one-way ANOVA results for clustered input regions for *Cck*_VTA_ (left) and *Crhr1*_VTA_ (right) groups (see Supplementary Table 1 for statistical values).

We visualized the brain-wide cellular inputs of *Cck*_VTA_ and *Crhr1*_VTA_ populations by assessing the mean cell density per voxels (**Figure 8C, left**), and their significant group differences (**Figure 8C, right**). An orthogonal analysis was also conducted on brain atlas-segmented counts with group mean statistical comparisons made on a region-by-region basis (**Figure 8D-8E**). The overall anatomy of identified input regions were largely consistent with previous input-mapping studies of VTA dopamine neurons (Watabe-Uchida 2012). For example, both dopamine populations received large proportions of their total input from the LH, periaqueductal gray (PAG), superior colliculus (SC), and dorsal raphe (DR) (**Figure 8C-8D**). However, we found significant differences in input density from multiple regions in the striatum, pallidum, amygdala, hippocampus, hypothalamus, midbrain, and hindbrain (**Figure 8C-8E**; Supplementary Table 1). In the striatum, the NAc shell predominantly contained *Cck*_VTA_ input neurons, whereas dorsal striatum predominantly contained *Crhr1*_VTA_ input neurons (**Figure 8C-8E**). Thus, *Cck*_VTA_ neurons have reciprocal connections with their NAc projection target, whereas *Crhr1*_VTA_ neurons do not. In pallidal areas, the ventral pallidum (VP) was primarily a source of *Cck*_VTA_ input neurons, whereas dorsal globus pallidus conversely contained inputs prefrentially to *Crhr1*_VTA_ neurons (**Figure 8C-8E**). In more posterior regions, we found that the central amygdala (CeA) predominantly contained *Crhr1*_VTA_ input neurons, whereas the hippocampus connected predominantly to *Cck*_VTA_ neurons (**Figure 8C-8E**). A variety of septal regions including the medial septal nucleus (MS), diagonal band (DB) and hypothalamic regions including medial preoptic (MPO) and lateral preoptic (LPO) areas, LH and posterior hypothalamic nuclei contained significantly more *Cck*_VTA_ input neurons (**Figure 8C-8E**). *Crhr1*_VTA_ neurons received significantly more input from zona incerta (ZI), and multiple midbrain and hindbrain regions (e.g., medial reticular formation (MRF), SC, PAG, substantia nigra pars reticular part, pontine reticular nucleus, and cuneiform nucleus) (**Figure 8C-8E**).

We additionally clustered brain-wide input differences between *Cck*_VTA_ and *Crhr1*_VTA_ neurons into canonical functional networks (Xu et al., 2022), and found that many were implicated in sensory and motor processing, sensory-motor integration, motivation and action, cognitive processing, and affective processing (**Figure 8E**; see **Materials and Methods**). Interestingly, we found that *Cck*_VTA_ neurons received preferential inputs from brain regions involved in motivation and action selection (**Figure 8E**), whereas *Crhr1*_VTA_ neurons received preferential inputs from brain regions involved in sensory-motor integration and affect (**Figure 8E**). Thus, the distinct input patterns of *Cck*_VTA_ and *Crhr1*_VTA_ subpopulations are anatomically widespread with functional relevance inherent across multiple brain systems. Taken together, these results suggest that *Cck*_VTA_ and *Crhr1*_VTA_ subpopulations integrate distinct types of information from upstream inputs and reveal a potential mechanism for a larger diversification of anatomically and functionally specialized inputs to VTA subpopulations.

## Discussion

Projection-defined VTA dopamine subpopulations and the VTA GABA population contribute to learning and motivation through distinct mechanisms. The *Cck*_VTA_ and *Crhr1*_VTA_ dopamine populations, but not the Vgat_VTA_ population, preferentially encode distinct features of prediction-error, reward anticipation, and action-cue information. By contrast, the Vgat_VTA_ population preferentially encodes reward retrieval and its response profile *in vivo* strongly contrasts those of VTA dopamine subpopulations. Both the neural activity dynamics and behavioral effects of neural manipulations of VTA dopamine subpopulations are consistent with distinct roles in learning and motivation. We also found that in addition to their NAc subregion synaptic projection target, *Crhr1*_VTA_ and *Cck*_VTA_ populations can be defined by their distinct electrophysiological properties, pattern of brain-wide monosynaptic input, and functional connectivity with inhibitory and disinhibitory VTA circuits.

### Distinct functions of activity in VTA subpopulations for reward learning and motivation

Theoretical models of learning in the brain propose that the dopaminergic modulatory input reaching the striatum contains a prediction-error signal to support reinforcement learning (Schultz, 1997; Doya 2000). Another foundational model suggests that prediction-error signaling cannot fully account for the function of midbrain dopamine in supporting goal-directed behavior. The incentive-sensitization hypothesis proposes that dopamine signaling in the striatum provides the link between the hedonic valuation of rewards and the assignment of value to stimuli and actions associated with them (Berridge and Robinson, 1998). Thus, prevailing hypotheses suggest that midbrain dopamine plays dual roles in both invigorating ongoing behavior (motivation) and guiding future behavior (learning) (Salamone and Correa, 2012). It has been proposed that dopamine release in the NAc core mediates value-based learning whereas dopamine release in the shell mediates motivational salience (Kelley 1999; Saddoris 2015). Consistent with this hypothesis, our previous results showed that dopamine neurons with projections to NAc core or shell subregions differentially regulate the acquisition, extinction, and maintenance of instrumental responding (Heymann et al., 2020). In addition, recent studies have demonstrated that VTA GABA neurons respond to rewarding and aversive stimuli and constrain the activity of VTA dopamine neurons during motivated behavior (Bouarab et al., 2019; Van Zessen et al., 2012; Eshel et al., 2015). However, few studies have directly compared the endogenous activity dynamics of VTA GABA neurons with VTA dopamine neurons. Our findings extend those of previous studies by revealing how the neural activity dynamics of projection-defined VTA dopamine subpopulations and the VTA GABA inhibitory population contribute to learning and motivation.

Here we tested and extended the hypothesis that reward-related associative learning is mediated by *Crhr1*_VTA_ neurons whereas *Cck*_VTA_ dopamine neurons are more involved in ongoing motivated responding. First, we found that the response profiles of projection-defined dopamine populations are broadly similar during a cued reinstatement task. This is consistent with existing evidence that dopamine neurons respond relatively uniformly (similar response profile to action, cue, and outcomes) during simple instrumental conditioning tasks (Engelhard et al.; 2019, Heymann et al., 2020; Schultz, 1998), although the temporal resolution of fiber photometry recording of GCaMP6m fluorescence may be insufficient for detecting task-relevant neural activity dynamics on subsecond timescales. Our recordings revealed patterns of brief, time-locked increases in activity in VTA dopamine populations during action, cue, and outcome periods. By contrast, activity in all VTA populations was sustained long after reward delivery. We additionally observed patterns of sustained activity on long timescales (10-s) prior to reward delivery preferentially in the *Cck*_VTA_ population. Gradual increases in dopamine neuron activity and release as animals perform goal-directed behaviors and approach rewards has been reported (Hamid et al., 2015; Wassum et al., 2012; Howe et al., 2013; Collins and Saunders, 2020; Farrell et al., 2022). Whether this pattern of activity reflects reward expectation, prediction-errors, sustained effort, or motivational engagement remains unclear. The sustained increase in activity we observed in the *Cck*_VTA_ population during the action-cue period prior to reward across acquisition sessions could reflect a reward anticipation signal or an overall increased level of motivational engagement at trial onset.

When reward probability or reward effort were dynamically varied during learning and motivation tasks we found heterogeneous responses with non-uniform encoding of prediction-error, behavioral variables, and reward anticipation across NAc core and NAc shell-projecting subpopulations. We found a striking difference in how VTA dopamine populations encode prediction-error and action-cue behavioral variables. During a random reward-omission instrumental task, *Crhr1*_VTA_ activity reflected a prediction-error-like signal, while *Cck*_VTA_ activity reflected a salience signal. Further, *Crhr1*_VTA_ neurons preferentially encoded cues whereas the *Cck*_VTA_ population similarly encoded both actions and cues. The observed positive modulation of dopamine neuron activity by current trial outcome and negative modulation by previous trial outcome support the notion that *Crhr1*_VTA_ neurons are more involved in prediction-error encoding compared to *Cck*_VTA_ neurons. By contrast, the *Cck*_VTA_ population preferentially encoded the trial initiation lever-press action compared to the non-trial active lever-press action, whereas the *Crhr1*_VTA_ population similarly encoded both the trial initiation action and non-trial action. Taken together, these results suggests that *Crhr1*_VTA_ neurons strongly encode reward-associated stimuli and integrate predictive information over time to drive future behavior, whereas *Cck*_VTA_ neurons strongly encode goal-directed actions and provide a salience signal to drive ongoing behavior.

We found that the VTA dopamine populations have striking differences in how they respond to increasing levels of effort during bouts of lever-pressing throughout a progressive ratio task. Previous studies showed that VTA dopamine activity is negatively correlated with effort and reward attainability (Gan et al., 2010; Hamid et al., 2015); however, whether this effect is uniform across distinct dopamine subpopulations remains unclear. Our observation that *Cck*_VTA_ neurons display sustained elevation in activity during lever-pressing early in progressive ratio sequences when rewards are more attainable but not later suggests that sustained activity in this population, but not in *Crhr1*_VTA_ neurons, reflects reward anticipation during ongoing motivated behavior. Together, these findings suggest that prediction-error-encoding, action-cue encoding, and reward-anticipation encoding are dissociable processes that involve dopamine projections to distinct subregions of the NAc. In contrast to previous studies (Mohebi et al., 2019), our results suggests that sustained cell body activity in a distinct subpopulation of VTA dopamine neurons during behavior contributes to ongoing motivated responding. Further, it was recently demonstrated that dopamine neurons in the medial VTA display sustained activity patterns during behavior and encode behavioral state, whereas anatomically distinct lateral VTA dopamine neurons display transient activity and encode behavioral rate-of-change (de Jong et al., 2024).

In addition to examining how projection-defined dopamine populations contribute to learning and motivation, we further sought to directly compare the endogenous activity dynamics of VTA dopaminergic and GABAergic populations. The most prominent difference in activity dynamics among VTA subpopulations was found in the VTA GABA neurons, which encoded reward retrieval much more than VTA dopamine populations. While dopamine populations showed phasic and tonic responses during instrumental responses and cues, the Vgat_VTA_ population responded selectively during reward outcome periods. When reward was available, Vgat_VTA_ responses were sustained throughout reward consumption. Unrewarded port entries, however, evoked smaller, brief increases in Vgat_VTA_ activity. Further, the Vgat_VTA_ population preferentially encoded rewarded port entry compared to unrewarded port entry but showed no preferential encoding of behavioral variables during the action-cue period. This reward outcome-dependent difference in activity is consistent with findings that Vgat_VTA_ neurons causally contribute to reward expectation-driven decreases in VTA dopamine neuron activity (Eshel et al., 2015). Taken together, these results suggest that VTA GABA neurons reflect reward outcome information and may suppress VTA dopamine activity during reward consumption. Given the observed distinct activity dynamics among VTA dopamine populations during periods of high VTA GABA activity, VTA dopamine subpopulations likely have distinct connectivity patterns with VTA GABA neurons and may receive inhibitory input from distinct subpopulations of VTA GABA neurons.

Our previous work showed that *Crhr1*_VTA_ and *Cck*_VTA_ neurons differentially facilitate the acquisition and extinction of a goal-directed instrumental response (Heymann et al., 2020). Additionally, recent work showed that NAc core and NAc shell-projecting dopamine populations differentially contribute to Pavlovian cue conditioning (Saunders et al., 2018). Given this, we hypothesized that these populations differentially contribute to cued reinstatement. In contrast to the similarity in activity dynamics observed during reinstatement across dopamine subpopulations, their causal roles in reinstatement behavior were distinctive. We found that the effect of cue-paired activation on cued reinstatement behavior depended on which dopamine subpopulation was targeted. Consistent with the idea that reward-predictive cues drive motivated behavior through *Crhr1*_VTA_ neurons, activation of *Crhr1*_VTA_ neurons during cue presentations robustly increased cued reinstatement. Given the observed activity dynamics of *Crhr1*_VTA_ and *Cck*_VTA_ neurons during cued reinstatement, we hypothesized that either or both of these populations contribute causally to cued reinstatement behavior. Indeed, we found that cue-paired inhibition of *Crhr1*_VTA_ neurons reduced cued reinstatement behavior. In contrast, inhibition of *Cck*_VTA_ neurons did not affect cued reinstatement. While photostimulation generated robust effects on behavior, photoinhibition had a more modest impact on reinstatement. This could be due to incomplete inhibition, but it is likely that local control of dopamine release and other inputs to the NAc including the prefrontal cortex, thalamus, hippocampus, and amygdala contribute to cued reinstatement. Additionally, a modest reduction in reinstatement could reflect a floor effect since mice continue to respond habitually following extinction. Further, modifying the timing of optical manipulation relative to action, cue, and reward could reveal more nuanced contributions of dopamine subpopulations to cued reinstatement.

Together, these results support the idea that similar patterns of dopamine neuron activity contribute to distinct aspects of behavior depending on NAc subregion target. One possible interpretation of these results is that NAc core-projecting *Crhr1*_VTA_ neurons are selectively involved in encoding the motivational value of reward-predictive stimuli to drive future behavior whereas the NAc shell-projecting *Cck*_VTA_ neurons are preferentially involved in encoding goal-directed actions and reward anticipation during ongoing motivated behavior (Kelley, 1999; Saunders et al., 2018).

### Heterogeneous intrinsic and circuit properties define distinct VTA dopamine subpopulations

The endogenous phasic responses of VTA dopamine neurons during behavior are likely largely driven by excitatory VTA inputs (Chergui et al., 1993; Zweifel et al., 2009). Consistent with the differences we observed in baseline neural activity during periods of task-related behavioral inactivity between *Cck*_VTA_ and *Crhr1*_VTA_ populations, our *ex vivo* electrophysiological results reveled that these dopamine subpopulations displayed distinct intrinsic properties that may regulate their endogenous activity. Specifically, the increased baseline excitability of *Cck*_VTA_ neurons could contribute to their sustained increase in activity observed during multiple behaviors and increased baseline calcium transient width and amplitude. Additionally, we observed a greater level of spontaneous postsynaptic inhibitory transmission in *Cck*_VTA_ neurons. The amplitude of spontaneous inhibitory postsynaptic currents is correlated with the strength of inhibitory synapses onto the postsynaptic neuron (Segal, 2010; Glasgow et al., 2019). This could be due to multiple mechanisms including differences in the number, location, or subunit composition of GABA receptors (Farrant et al., 2007; Dixon et al., 2014). Distinct inhibitory synaptic transmission mechanisms in dopamine subpopulations may underlie specific aspects of their different response profiles *in vivo*.

A prominent finding of our study is that Vgat_VTA_ neuron activation suppressed the activity of *Crhr1*_VTA_ neurons more strongly than that of *Cck*_VTA_ neurons. This finding is consistent with the observation that *Cck*_VTA_ and *Crhr1*_VTA_ neurons display distinct intrinsic properties. Previous input mapping studies have revealed that VTA dopamine and VTA GABA neurons receive inhibitory input from many of the same brain regions including LH, NAc, PAG, and DRN (Morales and Margolis, 2017; Yang et al., 2018). Recent studies have shown that specific GABAergic projections from the LH and NAc are important for behavioral activation and motivation, respectively (Lammel et al., 2012; Yang et al., 2018; Nieh et al., 2016). Interestingly, our functional optogenetic experiments revealed that NAc shell GABA activation disinhibits *Cck*_VTA_ neurons selectively. This dedicated NAc shell GABA-VTA GABA-*Cck*_VTA_ dopamine disinhibitory pathway could provide a mechanism by which information about motivational salience reflected in NAc shell MSN activity drives activity in *Cck*_VTA_ neurons during goal-directed actions. The observation that LH GABA provides a similarly strong disinhibitory drive onto both *Cck*_VTA_ and *Crhr1*_VTA_ neurons is consistent with previous findings that the LH GABA-VTA GABA-VTA dopamine circuit plays a broader role in positive reinforcement and behavioral activation across a wide range of motivated behaviors (Nieh et al., 2016). Further, these results are consistent with the hypothesis that VTA GABA neurons are heterogeneous in their afferent connectivity and in their synaptic connectivity with VTA dopamine subpopulations. Taken together, these findings support a model in which dopamine subpopulations are embedded in distinct inhibitory circuits which contributes to their distinct response profiles *in vivo*.

Given the observed heterogeneous activity dynamics and functional roles of *Crhr1*_VTA_ and *Cck*_VTA_ neurons during motivated behavior, we hypothesized that these populations receive distinct upstream monosynaptic inputs. Consistent with previous input mapping studies of VTA dopamine neurons, we identified ∼100 brain regions connected to *Crhr1*_VTA_ and *Cck*_VTA_ neurons, the majority of which were common to both cell types. Interestingly, *Crhr1*_VTA_ neurons receive a preferential density of inputs from brain regions involved in sensory-motor integration including dorsal striatum, globus pallidus, zona incerta, and superior colliculus. By contrast, *Cck*_VTA_ neurons receive a higher density of inputs from brain regions linked to motivation and action including NAc shell, ventral pallidum, and LH. Prior work has demonstrated that information about reward outcome, expectation, and prediction is distributed across VTA input neurons in regions such as the LH, dorsal and ventral striatum, rostromedial tegmental nucleus, and ventral pallidum among others (Tian e al., 2016). Further, activity in NAc shell projection neurons has been shown to reflect motivation during behavior (Castro et al., 2019; Floresco, 2015). Our results indicating differential monosynaptic input density from these regions to *Crhr1*_VTA_ and *Cck*_VTA_ populations suggest that non-uniform integration of reward-related information contributes to their distinct roles in learning and motivation.

## Materials and Methods

### Experimental Subject Details

Male and female mice, housed on a 12 h light/dark cycle, were used in all experiments performed during the light phase. All procedures were approved and performed under the guidelines of the Institutional Animal Care and Use Committee at the University of Washington. Mice between the ages of 2-6 months were used for all experiments. Vgat-Cre (*Slc32a1^tm2(cre)Lowl^/J*), DAT-Cre (B6.SJL-*Slc6a3^tm1.1(cre)Bkmn^*/J), and *Cck*-Cre (*Cck^tm1.1(cre)Zjh^*/J) were purchased from The Jackson Laboratory and bred in house. *Crhr1-*Cre mice were generated as previously described (Sanford et al., 2017).

### Cued Reinstatement Task

Mice were food-restricted to 85%-90% of their ad libitum weight and trained on a cued reinstatement task (adapted from Nugent et al., 2017; Soden et al., 2022) in Med-associates operant chambers with two retractable levers and a food port. During 1-2 pre-training sessions, mice were trained to press either lever for immediate delivery of a 20 mg sucrose pellet (Bio-Serv, F0071). All trials were followed by 3-s inter-trial interval. When mice earned 20 pellets within an hour of training, the session ended. In the acquisition task, mice were trained to press the active lever which initiated a 3-s delay period followed by a 3-s cue period (active lever cue light and 2.9 kHz continuous tone). At the end of the cue period, a sucrose pellet was delivered into the food port. At the cue period onset, the box house light was extinguished until the end of the 12.5-s inter-trial interval period. In the extinction task, mice were free to press either lever but all cues and rewards were omitted. In the reinstatement task, 5 non-contingent cue presentations were delivered every 2 min for 10 min during a pre-session period in which the levers were not extended into the box. The pre-session was immediately followed by a 1 h session identical to the acquisition phase except without delivery of food reinforcers.

### Random Reward-Omission Task

In the random reward omission task, an active lever press initiated a 3-s delay period followed by a 3-s cue period (active lever cue light and 2.9 kHz continuous tone). Then, at cue offset mice received a food reward on ∼50% of trials. Rewarded and unrewarded trials were randomly interspersed throughout the session so that upcoming trial outcomes were unpredictable. At the cue period onset, the box house light was extinguished until the end of the 12.5-s inter-trial interval period regardless of trial outcome type.

### Progressive Ratio Task

Food-restricted mice were trained on a fixed-ratio (FR1) schedule of reinforcement for food reward during three daily one hour sessions in which each active lever press resulted in delivery of a sucrose pellet after a 3-s delay. Then, mice underwent a single session with a progressive ratio schedule of reinforcement in which the number of active lever presses required for each food reinforcer is increased after the completion of each ratio (i.e. 1, 2, 4, 6, 9, 13…) following a pseudo-exponential function. If no lever press responses were made within a 3 m time period or 120 m elapsed, the session ended.

### Surgery

Mice (6-8 weeks) were anesthetized with isoflurane (1.5 – 4%) and head-fixed for stereotaxic (David Kopf Instruments) survival surgery. Stereotactic coordinates were standardized relative to Bregma and Lambda distance and an injection syringe was used to inject 0.5 µL of virus at a rate of 0.25 µL/min. Mice recovered from surgery for at least two weeks prior to behavioral testing.

#### Fiber photometry

For fiber photometry behavior experiments, 0.5 µL of AAV1-DIO-GCaMP6m was injected unilaterally into the VTA (A/P: -3.25 mm, M/L: -0.5 mm, D/V: -4.5 mm). Following the virus injection, a fiber optic cannula (400 µm) was implanted 0.5 mm above the VTA.

#### Optogenetics

For optogenetics activation behavior experiments, 0.5 µL of AAV1-DIO-ChR2-YFP was injected unilaterally into the VTA (A/P: -3.25 mm, M/L: -0.5 mm, D/V: -4.5 mm). Following the virus injection, a fiber optic cannula (200 µm) was implanted 0.5 mm above the VTA.

For optogenetic inhibition behavior experiments, 0.5 µL of AAV1-DIO-JAWS-GFP was injected bilaterally into the VTA (A/P: -3.25 mm, M/L: ±0.5 mm, D/V: -4.5 mm). Following the virus injection, fiber optic cannulae (200 µm) were implanted 0.5 mm above the VTA bilaterally. The fiber optic cannula was implanted at an angle of 10 degrees on one side.

For optogenetic activation and inhibition experiments, control mice were injected with AAV1-DIO-EGFP-KASH.

#### Dual optogenetic stimulation and fiber photometry recording

For dual stimulation and recording experiments of VTA GABA neurons and VTA dopamine subpopulations, 0.5 µL of equal parts AAV1-DIO-Chrimson-TdTomato and AAV1-DIO-GCaMP6m was injected unilaterally in to the VTA (A/P: -3.25 mm, M/L: -0.5 mm, D/V: -4.5 mm). Following the virus injection, a fiber optic cannula (400 µm) was implanted 0.5 mm above the VTA.

For dual stimulation and recording experiments of LH GABA neurons and VTA dopamine subpopulations, 0.5 µL of AAV1-DIO-Chrimson-TdTomato was injected unilaterally in to the LH (A/P: -1.35 mm, M/L: -1.0 mm, D/V: -5.0 mm) and AAV1-DIO-GCaMP6m was injected unilaterally in the VTA (A/P: -3.25 mm, M/L: ±0.5 mm, D/V: -4.5 mm). Following the virus injection, a fiber optic cannula (200 um) was implanted 0.5 mm above the LH at an angle of 5 degrees and a fiber optic cannula (400 µm) was implanted above the VTA.

For dual stimulation and recording experiments of NAc medial shell GABA neurons and VTA dopamine subpopulations, 0.5 µL of AAV1-DIO-Chrimson-TdTomato was injected unilaterally in to the NAc medial shell (A/P: 1.25 mm, M/L: -0.6 mm, D/V: -4.6 mm) and AAV1-DIO-GCaMP6m was injected unilaterally in the VTA (A/P: -3.25 mm, M/L: ±0.5 mm, D/V: -4.5 mm). Following the virus injection, a fiber optic cannula (200 µm) was implanted 0.5 mm above the NAc medial shell and a fiber optic cannula (400 µm) was implanted above the VTA.

#### Ex vivo slice electrophysiology

For electrophysiology experiments, 0.5 µL of AAV1-DIO-YFP was injected bilaterally into the VTA (A/P: -3.25 mm, M/L: -0.5 mm, D/V: -4.5 mm).

#### Rabies retrograde tracing

For the retrograde tracing experiment, 0.5 µL of AAV-syn-DIO-TC66T-2A-eGFP-2A-oG was injected bilaterally into the VTA (A/P: -3.45 mm, M/L: ±0.5 mm, D/V: -4.5 mm). 14 days later, 0.5 µL of EnvA-SADΔG-RV-DsRed was injected bilaterally into the VTA (A/P: -3.45 mm, M/L: ± 0.5 mm, D/V: -4.5 mm).

### Fiber photometry

GCaMP6m fluorescence was recorded through an implanted optic fiber connected to a path cord (Doric Lenses) while mice performed behavioral tasks. An LED driver (Doric Lenses) was used to control two LEDs for excitation of GCaMP6m. A 465-nm LED (light intensity: 30-40 µW, sinusoidal frequency modulation: 531 Hz) was used to excite GCaMP6m for calcium-dependent fluorescence. A 405-nm LED (light intensity: 30-40 µW, sinusoidal frequency modulation: 211 Hz) was used to excite GCaMP6m for calcium-independent fluorescence. Light intensity was measured at the optic fiber tip. GCaMP6m fluorescence was collected through the same optic fiber using a photoreceiver (Doric Lenses) and recorded using a Tucker Davis Technologies (TDT) real-time processor (RZ5 BioAmp) at 1017.25 Hz sampling frequency. Task event timestamps were simultaneously registered as TTL pulses by the MedAssociates system and synchronously delivered to the TDT system through a custom interface.

### Fiber photometry preprocessing

The 405-nm and 465-nm signals were demodulated and downsampled to 50 Hz using a moving average. The downsampled 405-nm control signal was fit to the 465-nm GCaMP6m signal using the ‘np.polyfit’ function in Python with a degree of 1 to obtain a fitted control signal. The fitted control signal was subtracted from the downsampled 465-nm GCaMP6m signal to correct for calcium-independent fluorescence changes. Then, an exponential curve was fit to the corrected 465-nm signal and subtracted to correct for slow baseline drift due to photobleaching. The baseline-corrected 465-nm GCaMP6m signal for the entire recording session (ΔF/F) was z-scored relative to the mean and standard deviation of the entire session trace to allow for comparison across individual recording sessions and individual mice. The session z-score was defined as ΔF/F(t) - mean(ΔF/F(t)) / std(ΔF/F(t)). The preprocessed photometry signal was aligned to task events and mean z-score ± SEM was calculated for behavioral epochs of interest. Lever pressing and port entry bouts were classified in either 10-s or 30-s windows aligned to session event times and the first event time per bout (bout onset) was aligned to the photometry signal. Baseline calcium transient analysis was restricted to time periods in which no behavioral event timestamps (lever-press, port entry) were recorded for at least 60-s during the final extinction session of the cued reinstatement task. This resulted in ∼20-30 min of baseline recording time per mouse. Transients were identified using the Python ‘scipy.signal.find_peaks’ function.

### Optogenetics

For behavioral experiments with optogenetics, mice underwent the pretraining, acquisition, and extinction phases of the cued reinstatement task without laser stimulation. For photostimulation experiments, mice received blue light (3-s, 20 Hz, 5-ms pulse width, ∼10 mW light power) unilaterally in the VTA 3-s following active lever press throughout the control stimulation session following the last extinction session. Throughout the cued reinstatement session, mice received blue light (3-s, 20 Hz, 5-ms pulse width, ∼10 mW) unilaterally in the VTA during cue presentation. For photoinhibition experiments, mice received red light (2-s constant square pulse terminated with a 1-s linear ramp-down, ∼5-10 mW light power) unilaterally in the VTA during cue presentation throughout the cued reinstatement session.

For photostimulation of VTA GABAergic neurons in the VTA, mice were placed in an operant box and received red light (1-s or 3-s, 5 - 40 Hz, 5-ms pulse width, ∼10 mW light power) every 60-s for 80 trials across two sessions. Red light was delivered through the same optic fiber used for fiber photometry recording.

For stimulation of LH or Nac medial shell GABAergic neurons, mice received red light (1-s or 3- s, 5 - 40 Hz, 5-ms pulse width, ∼10 mW light power) every 60-s for 80 trials across two sessions. Red light was delivered through an optic fiber above the LH or Nac medial shell.

### Ex vivo electrophysiology

Horizontal VTA sections (200 µm) were prepared in a NMDG cutting solution (92 NMDG mM, 2.5 KCl mM, 1.25 NaH_2_PO_4_ mM, 30 NaHCO_3_ mM, 20 HEPES mM, 25 glucose mM, 2 thiourea mM, 5 Na-ascorbate mM, 3 Na-pyruvate mM, 0.5 CaCl_2_ mM, 10 MgSO_4_ mM, pH 7.3–7.4). Then, sections were incubated for ∼12 min in the same solution at 32°C in a water bath. Slices were then transferred to a HEPES-aCSF solution (92 NaCl mM, 2.5 KCl mM, 1.25 mM NaH_2_PO_4_ mM, 30 NaHCO_3_ mM, 20 HEPES mM, 25 glucose mM, 2 thiourea mM, 5 Na-ascorbate mM, 3 Na-pyruvate mM, 2 CaCl_2_ mM, 2 MgSO_4_ mM) at room temperature. Slices recovered for an additional 60 min.

Whole-cell, patch-clamp recordings were acquired using the Axopatch 700B amplifier (Molecular Devices) at a sampling frequency of 10 kHz and filtering at 1 kHz. eYFP-positive cells were visualized using fluorescence for recording with electrodes at 3–5 MΩ. Excitability recordings were made in an aCSF solution (126 NaCl mM, 2.5 KCl mM, 1.2 NaH_2_PO_4_ mM, 1.2 MgCl_2_ mM, 11 D-glucose mM, 18 NaHCO_3_ mM, 2.4 CaCl_2_ mM) at 32°C and for sIPSC recordings the solution was supplemented with 2mM of kynurenic acid. The aCSF solution was perfused over slices at ∼2 ml/min.

To determine excitability, cells were patched with electrodes were filled with an internal solution (in mM: 130 potassium gluconate, 10 HEPES, 5 NaCl, 1 EGTA, 5 Mg-ATP, 0.5 Na-GTP, pH 7.3, 280 mOsm). Current-voltage curves were generated by recording in current-clamp mode and injecting steps of current (0-80-pA, 10-pA steps, 1-s) (both at resting membrane potential and at -60 mV). For sIPSCs, cells were patched with electrodes were filled with an internal solution (in mM: 135 KCl, 12 NaCl, 0.5 EGTA, 10 HEPES, 2.5 Mg-ATP, 0.25 sodium GTP, pH 7.3, 280 mOsm) and held at -60 mV. Recordings were analyzed using Clampfit (Molecular Devices). Excitability was calculated as the total number of events during each current step detected using the Event Detection function in Clampfit (50-ms before and 150-ms after action potential peak). The first action potential evoked from a 20-pA current injection while holding at -60mV were detected using the Event Detection threshold function in Clampfit. Hyperpolarization-induced rebound activity was assessed by injecting a hyperpolarizing step of current (-120-pA, 1-s). sIPSCS were detected using the Event Detection template function in Clampfit to determine frequency and amplitude.

### Retrograde tracing

Mice were perfused and brains were collected for imaging (see Immunohistochemistry) nine days following rabies virus injection (see Surgery). Then we optically cleared whole-brain samples using the SmartClear full active pipeline protocol (LifeCanvas Technologies, v5.05) for aqueous-based brain clearing and mounting as described recently (Szelenyi et al, 2023). Native fluorescent signals from aqueous-based cleared brains were imaged horizontally using the SmartSPIM LSFM (LifeCanvas Technologies) at 4 µm near-isotropic pixel resolution in 2 channels: 488-nm for registration signal and helper virus-infected cells, and 563-nm for rabies virus-infected inputs cells. Laser power and acquisition settings were held constant for all genetic groups and their individuals. Cleared whole-brain samples were then placed in phosphate-buffered saline (PBS) for at least two 24 hour washes at room temperature. Whole-brain samples were then mounted in 4% agarose and sectioned into 50 µm sections using a vibratome for posthoc immunostaining of free-floating sections (see Immunohistochemistry and image analysis).

### Immunohistochemistry and image analysis

Mice were deeply anesthetized using Beuthanasia and transcardially perfused with phosphate-buffered saline (PBS), followed by 4% paraformaldehyde (PFA) in PBS. Brains were placed in 4% PFA overnight, then transferred to 30% sucrose in PBS solution at 4 ° C for at least 24 hours. Brains were then sectioned into 30-40 µm sections using a cryostat and placed in PBS at 4 ° C. Brain sections were then stained to validate virus expression. Free-floating sections were placed in blocking buffer (3% normal donkey serum and 0.3% Triton X-100 in PBS) for 30 min. For enhancement of GCaMP6m, ChR2-YFP, JAWS-GFP, TC66T-2A-eGFP, and eYFP, sections were incubated in primary antibody (Chicken-GFP, 1:6000 dilution, ABCAM) overnight at 4 ° C. For enhancement of Chrimson-TdTomato, sections were incubated in primary antibody (Rabbit-DsRed, 1:1000 dilution, Takara) overnight at 4 ° C. For detection of tyrosine hydroxylase sections were incubated in primary antibody (Rabbit-Tyrosine Hydroxylase, 1:1000 dilution, Millipore Sigma) overnight at 4 ° C. Sections were then placed in PBS for three ten minute washes, and incubated in secondary antibody (Alexa Fluor 488 AffiniPure Donkey Anti-Chicken or Cy3 Donkey Anti-Rabbit, Jackson ImmunoResearch) for 1 hour at room temperature. Then, following three ten minute PBS washes, sections were mounted using a mounting medium (DAPI Fluoromount-G, Southern Biotech) and coverslipped. Images were taken using a KEYENCE BZ-X fluorescence microscope (KEYENCE). Histology sections from mice with optic fiber implants were used to identify optic fiber tip locations. One section containing the optic fiber tip location per mouse was used for the cell count and fluorescence intensity quantification of GCaMP-positive cells for the mice used in fiber photometry experiments during behavior. A custom CellProfiler 4.1.3 pipeline was used for quantification of GFP-positive cells for quantification of GCaMP-positive cells in the photometry experiment and GFP-positive starter cells in the rabies tracing experiment. Fluorescence intensity was normalized to the maximum intensity per section.

## Quantification and Statistical Analysis

### Statistical Analysis

All data are expressed as mean ± SEM. Statistical analyses were performed using GraphPad Prism, Python, and MATLAB. All statistical tests were two-tailed. Sample sizes were not predetermined using statistical methods. For data from two groups, the paired *t*-test, unpaired *t*-test, and Mann-Whitney *U* test were used where appropriate. For data from three or more groups, one-way ANOVA and one-way repeated-measures ANOVA followed by multiple-comparisons tests (Tukey’s multiple comparisons test, Bonferroni’s multiple comparisons test, two-stage linear step-up procedure of Bejamini, Krieger and Yekutieli) were used to determine any statistically significant differences between groups. For data from three or more groups and across multiple conditions, two-way ANOVA and two-way repeated-measures ANOVA followed by multiple-comparisons tests (Šídák’s multiple comparisons test, two-stage linear step-up procedure of Bejamini, Krieger and Yekutieli) were used where appropriate. For correlation analysis, the Pearson correlation coefficient was used. For all tests, a significance threshold of 0.05 was used. See supplementary Table 1 for detailed statistical results.

### Linear Encoding Model

To quantify the relative contribution of task variables to neural activity, we used a linear encoding model (adapted from Engelhard et al., 2019 and Parker et al., 2022). Multiple linear regression was used to predict the photometry signal for a given mouse using task behavioral variables as predictors. The task predictor set consisted of a matrix of 10 behavior event types (event times of trial initiation lever press, non-trial active lever press, non-trial inactive lever press, cue onset, reward outcome, omission outcome, rewarded port entry, unrewarded port entry, non-trial port entry, trial-reset house light cue). Each event type time series was convolved with a set of cubic splines that span several seconds after the event and three seconds before an action event type. A longer set of cubic splines was used for rewarded and unrewarded trial outcome events to reflect the longer trial outcome neural responses. The encoding model is expressed as y = βX + ε, where y is the GCaMP6m fluorescence for a given mouse during the random reward omission task, X is the set of event predictors generated from the convolution of event times with the cubic spline set, and β is the set of weights learned from the regression.

We first fit the full version of the encoding model using the ‘fitglm’ function in MATLAB with threefold cross-validation to generate R^2^ for the full model. We then compared the model fit when each of the task predictors were removed to that of the full model to quantify the relative contribution of individual behavioral variables. An individual task predictor contribution was defined as the reduction in explained variance ΔR^2^ when that task predictor was removed from the model (1 - R^2^ partial/ R^2^ full). The relative contribution of an individual task predictor was defined as a fraction of the predictor contribution over the full model predictor contributions.

### Reward Outcome History RPE Analysis

We used linear regression to predict neural activity following trial outcome using trial outcome information from the current and five previous trials (adapted from Bayer and Glimcher, 2005). The z-scored GCaMP6m signal during a 2-s time bin prior to trial initiation was subtracted from the z-scored GCaMP6m signal from 0 to 20-s following trial outcome for each trial during the random reward omission task. Current and previous trial outcomes were labeled 0 for omission and 1 for reward. Multiple linear regression was used to generate weights corresponding to the contribution of each of the current and five previous trial outcomes to the neural activity following the current trial outcome. The model is expressed as y(t) = β_0_ + β_1_T_out_(t) + β_2_T_out_(t-1) + … + β_6_T_out_(t-5), where y(t) is the mean z-scored GCaMP6m signal from 0 to 20-s on trial t, T_out_(t) is the trial outcome, and β_i_ is the regression coefficient for trial T_out_(t-n). The regression coefficients for each trial lag were generated with the ‘OLS’ function from the ‘linear_model’ module in the Python ‘statsmodels’ package.

### Whole-brain image processing and quantification

We used ImageJ software to crop whole-brain image stacks, transform from the horizontal to coronal plane, and export images as TIFF files for whole-brain analysis. Brains were registered to the Unified brain atlas and segmented DsRed cell counts were partitioned into regions with Unified atlas labels (Chon et al., 2019). For brain atlas registration and automated cell detection a previously described and modified ClearMap analysis pipeline (Renier et al., 2016; Madangopal et al., 2022) was used with minor adjustments. Accordingly, cell segmentation was automated using the Spot Detection function. The number of DsRed-positive input neurons per brain region was normalized to the total volume of the brain region from the reference atlas to calculate cell density per region (cells/mm^3^). 3D renders of input cell location in atlas space were generated using the ‘Wholebrain’ (Furth et al., 2018) and ‘SMART’ (Jin et al., 2022) packages in R.

---

**Figure S1.**
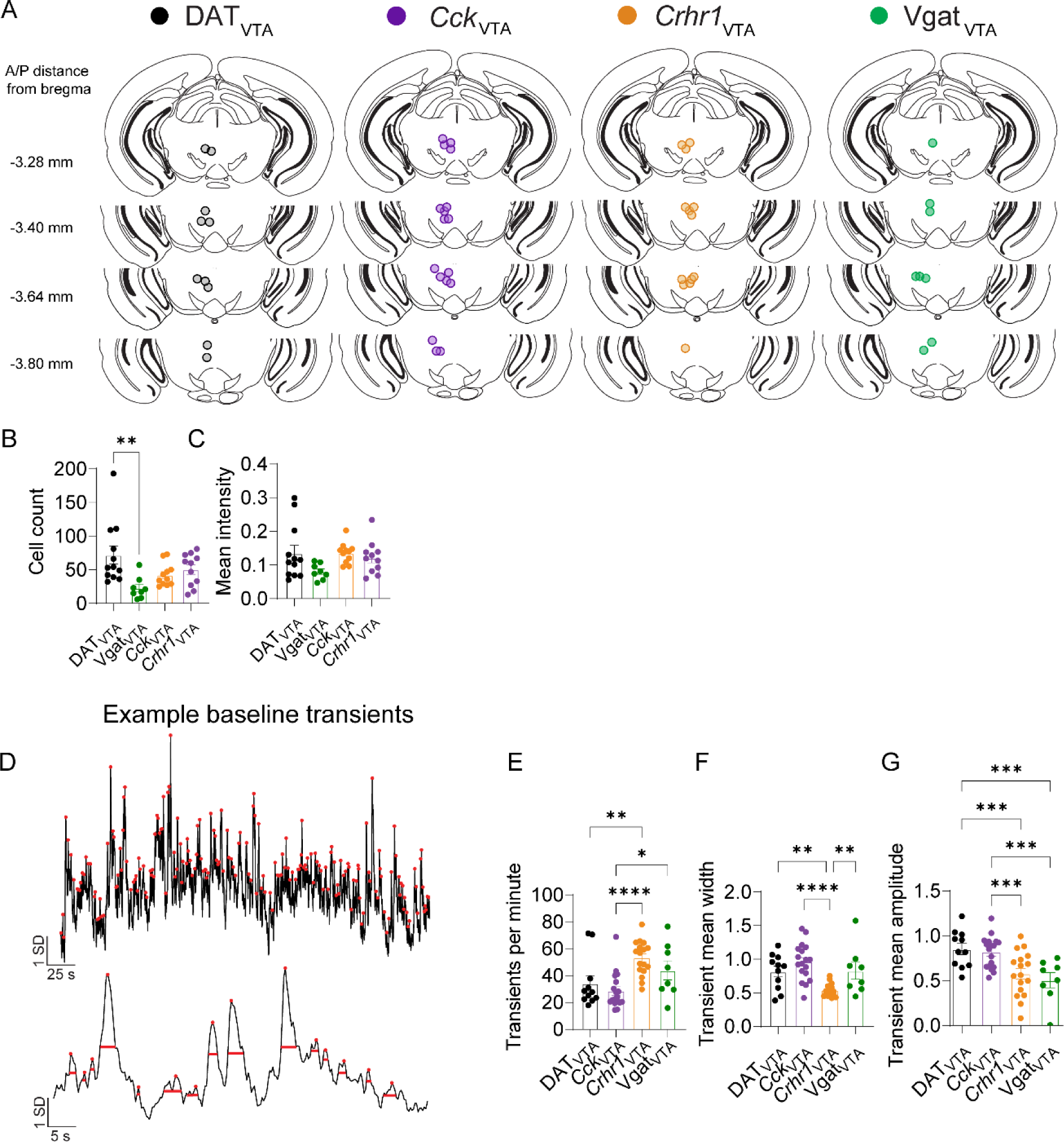
Summary of optic fiber tip locations for fiber photometry recordings, GCaMP-labeled cell counts and intensity measurements, baseline calcium transient analysis related to Figures 1-4. (A) Photometry recording fiber locations for DATVTA (n = 10 mice), *Cck*_VTA_ (n = 17 mice), *Crhr1*_VTA_ (n = 13 mice), and Vgat_VTA_ (n = 8 mice). Circles indicate the fiber tip location from individual mice. (B) Mean number of GFP-positive cells in the VTA in the histology section containing the optic fiber tip location for DATVTA (n = 10 mice), *Cck*_VTA_ (n = 17 mice), *Crhr1*_VTA_ (n = 13 mice), and Vgat_VTA_ (n = 8 mice). (C) Mean GFP fluorescence intensity in the VTA in the histology section containing the optic fiber tip location for DATVTA (n = 10 mice), *Cck*_VTA_ (n = 17 mice), *Crhr1*_VTA_ (n = 13 mice), and Vgat_VTA_ (n = 8 mice). (D) Example z-scored GCaMP fluorescence trace showing transient peak classifications (red circles) and transient width measurements (red bars) during baseline photometry recordings. (E) Mean transients per minute for *Cck*_VTA_ (n = 18 mice), *Crhr1*_VTA_ (n = 17 mice), DATVTA (n = 11 mice), Vgat_VTA_ (n = 8 mice) groups. (F) Mean transient width for *Cck*_VTA_ (n = 18 mice), *Crhr1*_VTA_ (n = 17 mice), DATVTA (n = 11 mice), Vgat_VTA_ (n = 8 mice) groups. (G) Mean transient amplitude for *Cck*_VTA_ (n = 18 mice), *Crhr1*_VTA_ (n = 17 mice), DATVTA (n = 11 mice), VgatVTA (n = 8 mice) groups. Bars and error bars indicate mean ± SEM across mice (see Supplementary Table 1 for statistical values).

**Figure S2.**
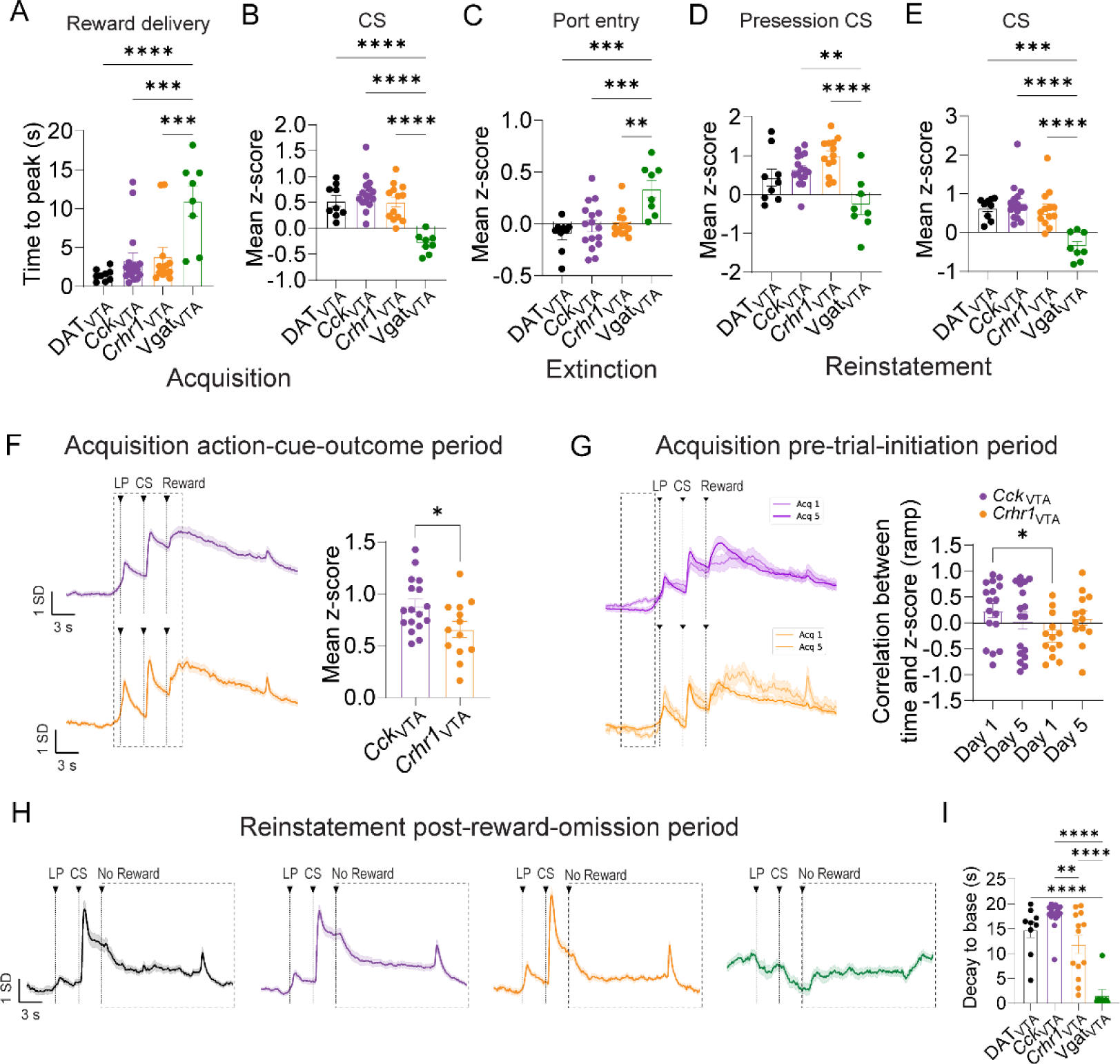
Further characterization of response profiles, related to Figure 1. (A) Average latency to peak of z-scored GCaMP fluorescence following reward delivery during acquisition phase of cued reinstatement for DAT_VTA_ (n = 9 mice), *Cck*_VTA_ (n = 16 mice), *Crhr1*_VTA_ (n 13 = mice), Vgat_VTA_ (n = 8 mice) groups. (B) Average z-scored GCaMP fluorescence during CS presentation period following reward delivery during acquisition phase of cued reinstatement for DAT_VTA_ (n = 9 mice), *Cck*_VTA_ (n = 16 mice), *Crhr1*_VTA_ (n 13 = mice), Vgat_VTA_ (n = 8 mice) groups. (C) Average z-scored GCaMP fluorescence during port entry period during extinction phase of cued reinstatement for DAT_VTA_ (n = 9 mice), *Cck*_VTA_ (n = 16 mice), *Crhr1*_VTA_ (n 13 = mice), Vgat_VTA_ (n = 8 mice) groups. (D) Average z-scored GCaMP fluorescence during CS presentation periods during reinstatement presession for DAT_VTA_ (n = 9 mice), *Cck*_VTA_ (n = 16 mice), *Crhr1*_VTA_ (n 13 = mice), Vgat_VTA_ (n = 8 mice) groups. (E) Average z-scored GCaMP fluorescence during CS presentation periods during reinstatement session for DAT_VTA_ (n = 9 mice), *Cck*_VTA_ (n = 16 mice), *Crhr1*_VTA_ (n 13 = mice), Vgat_VTA_ (n = 8 mice) groups. (F) Z-scored GCaMP fluorescence aligned to action-cue period during acquisition phase of cued reinstatement in *Cck*_VTA_ (n = 16 mice) and *Crhr1*_VTA_ (n = 13 mice) groups. Dotted rectangle indicates mean z-score analysis epoch. (G) Z-scored GCaMP fluorescence aligned to trial initiation LP on the first (lighter shade) and last (darker shade) session of acquisition in *Cck*_VTA_ (n = 16 mice) and *Crhr1*_VTA_ (n = 13 mice) groups (left). Pearson’s correlation coefficient per mouse between time and z-scored GCaMP signal during the time period prior to trial initiation LP on the first and last session of acquisition for *Cck*_VTA_ (n = 16 mice) and *Crhr1*_VTA_ (n = 13 mice) groups (right). Dotted rectangle indicates analysis epoch. Bars and error bars indicate mean ± SEM across mice. (H) Z-scored GCaMP fluorescence aligned to reward omission period during reinstatement phase of cued reinstatement in *Cck*_VTA_ (n = 16 mice) and *Crhr1*_VTA_ (n = 13 mice) groups. Dotted rectangle indicates mean z-score analysis epoch. (I) Average latency to minimum GCaMP fluorescence during reward omission period during reinstatement session for *Cck*_VTA_ (n = 16 mice) and *Crhr1*_VTA_ (n = 13 mice) groups. Bars and error bars indicate mean ± SEM across mice (see Supplementary Table 1 for statistical values).

**Figure S3.**
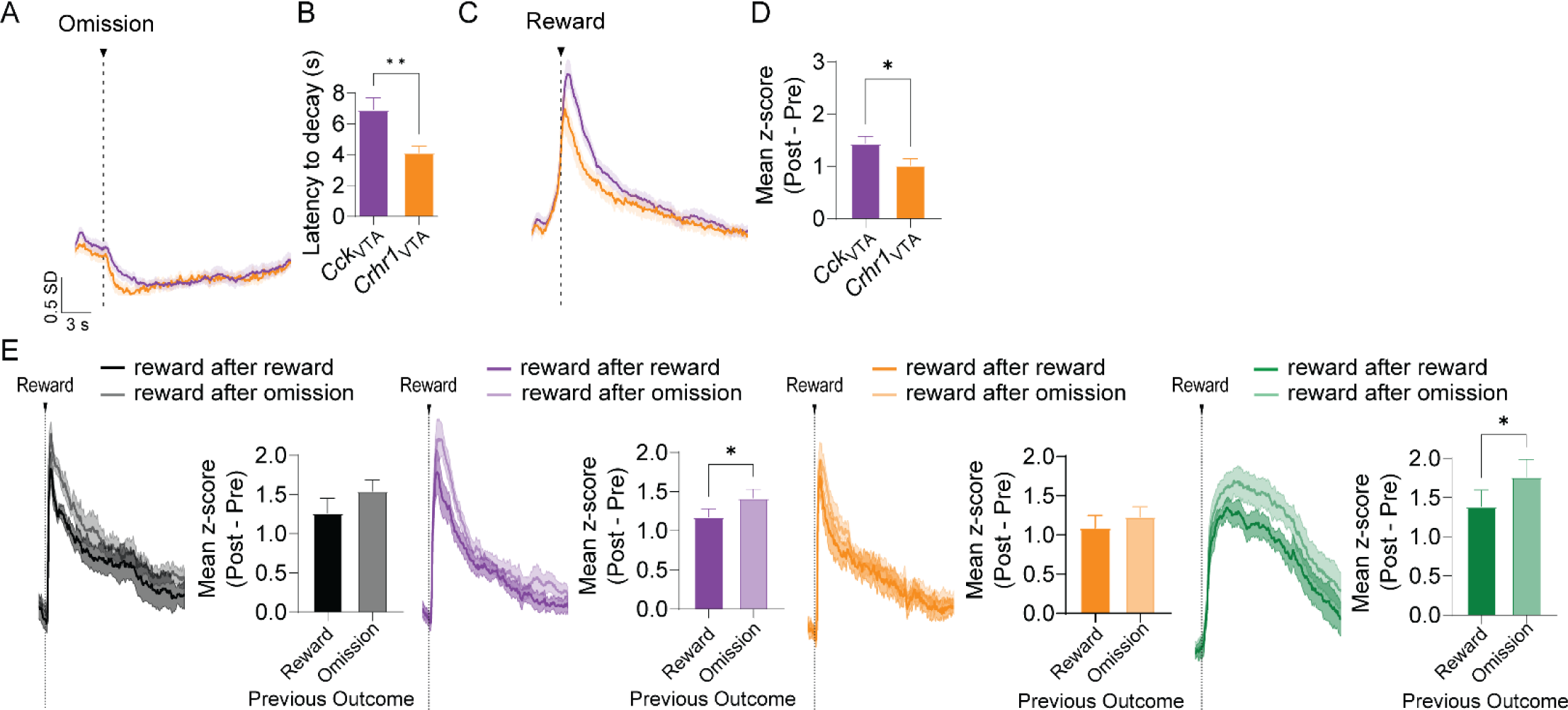
Further characterization of response profiles, related to Figure 2. (A) Z-scored GCaMP fluorescence following reward omission during random reward omission for DAT_VTA_ (n = 7 mice), *Cck*_VTA_ (n = 12 mice), and *Crhr1*_VTA_ (n = 13 mice), and Vgat_VTA_ (n = 8 mice) groups. (B) Average latency to trough of z-scored GCaMP fluorescence following reward during random reward omission for DAT_VTA_ (n = 7 mice), *Cck*_VTA_ (n = 12 mice), and *Crhr1*_VTA_ (n = 13 mice), and Vgat_VTA_ (n = 8 mice) groups. (C) Z-scored GCaMP fluorescence and average z-scored GCaMP fluorescence during reward omission periods shaded according to previous trial outcome type for DAT_VTA_ (n = 7 mice), *Cck*_VTA_ (n = 12 mice), and *Crhr1*_VTA_ (n = 13 mice), and Vgat_VTA_ groups (n = 8 mice). (D) Average baselin-subtracted z-scored GCaMP fluorescence following reward during random reward omission for DAT_VTA_ (n = 7 mice), *Cck*_VTA_ (n = 12 mice), and *Crhr1*_VTA_ (n = 13 mice), and Vgat_VTA_ (n = 8 mice) groups. (E) Z-scored GCaMP fluorescence and average z-scored GCaMP fluorescence during reward omission periods shaded according to previous trial outcome type for DAT_VTA_ (n = 7 mice), *Cck*_VTA_ (n = 12 mice), and *Crhr1*_VTA_ (n = 13 mice), and Vgat_VTA_ groups (n = 8 mice, see Supplementary Table 1 for statistical values).

**Figure S4.**
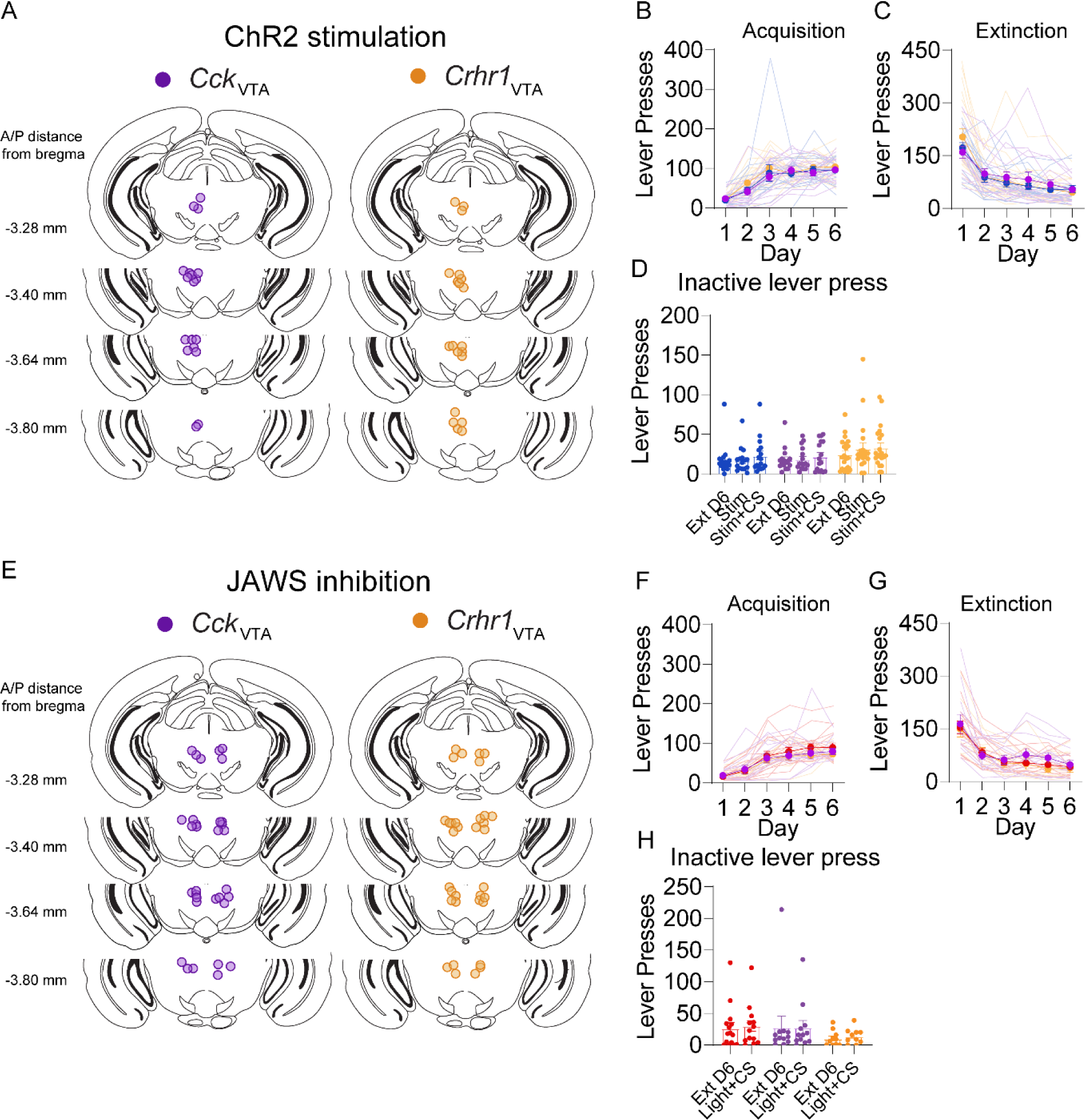
Summary of optic fiber tip locations for optogenetic manipulations, behavioral performance of mice during optogenetic cued reinstatement task, related to Figure 5. (A) Optic fiber tip locations in *Cck*_VTA_ (n = 15 mice) and *Crhr1*_VTA_ (n = 21 mice) groups for the ChR2 stimulation experiment. Circles indicate the fiber tip location from individual mice. (B) Mean number of active lever-presses across mice during acquisition sessions in control, *Cck*_VTA_, and *Crhr1*_VTA_ groups (n = 11-21 mice, error bars represent SEM). (C) Mean number of active lever-presses across mice during extinction sessions in control, *Cck*_VTA_, and *Crhr1*_VTA_ groups (n = 11-21 mice, error bars represent SEM). (D) Mean number of inactive lever-presses across mice during final extinction session, Stim session, and Stim + CS session in control, *Cck*_VTA_, and *Crhr1*_VTA_ groups (n = 11-21 mice, error bars represent SEM). (E) *Cck*_VTA_ optic fiber locations in *Cck*_VTA_ (n = 13 mice) and *Crhr1*_VTA_ (n = 14 mice) groups for the JAWS inhibition experiment. Circles indicate the fiber tip location from individual mice. (F) Mean number of active lever-presses across mice during acquisition sessions in control, *Cck*_VTA_, and *Crhr1*_VTA_ groups (n = 12-14 mice, error bars represent SEM). (G) Mean number of active lever-presses across mice during extinction sessions in control, *Cck*_VTA_, and *Crhr1*_VTA_ groups (n = 12-14 mice, error bars represent SEM). (H) Mean number of inactive lever-presses across mice during final extinction session, Stim session, and Stim + CS session in control, *Cck*_VTA_, and *Crhr1*_VTA_ groups (n = 12-14 mice, error bars represent SEM, see Supplementary Table 1 for statistical values).

**Figure S5.**
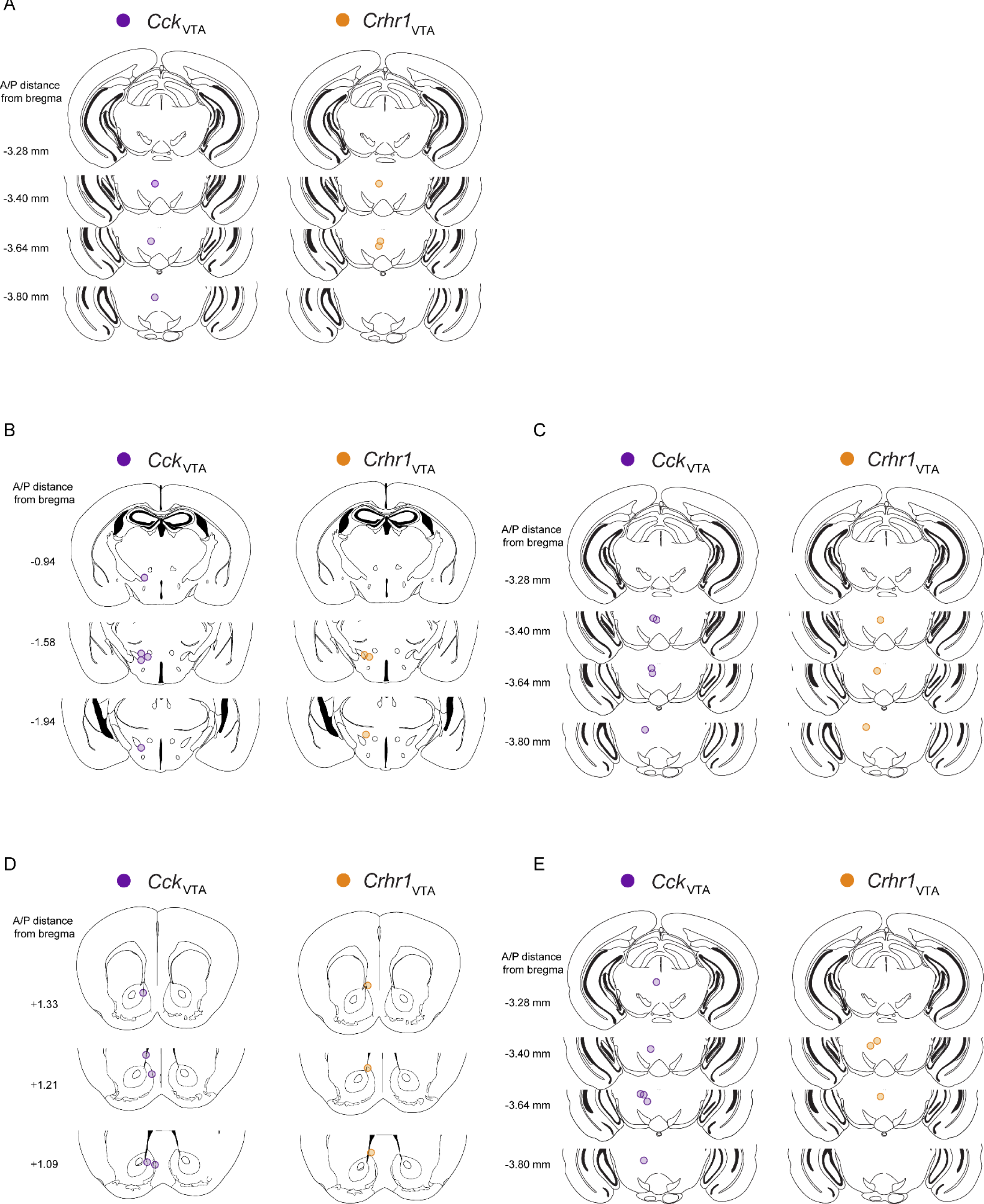
Summary of optic fiber tip locations for dual optogenetic stimulation and fiber photometry recordings. (A) Optic fiber tip locations in the VTA *Cck*_VTA_ (n = 3 mice) and *Crhr1*_VTA_ (n = 3 mice) groups for the Vgat_VTA_ stimulation with dual photometry experiment. Circles indicate the fiber tip location from individual mice. (B) Optic fiber tip locations in the LH for *Cck*_VTA_ (n = 5 mice) and *Crhr1*_VTA_ (n = 3 mice) groups for the LH GABA stimulation with dual photometry experiment. (C) Optic fiber tip locations in the VTA for *Cck*_VTA_ (n = 5 mice) and *Crhr1*_VTA_ (n = 3 mice) groups for the LH GABA stimulation with dual photometry experiment. (D) Optic fiber tip locations in the NAc mshell for *Cck*_VTA_ (n = 5 mice) and *Crhr1*_VTA_ (n = 3 mice) groups for the NAc mshell GABA stimulation with dual photometry experiment. Optic fiber tip locations in the VTA for *Cck*_VTA_ (n = 5 mice) and *Crhr1*_VTA_ (n = 3 mice) groups for the NAc mshell GABA stimulation with dual photometry experiment (see Supplementary Table 1 for statistical values).

**Figure S6.**
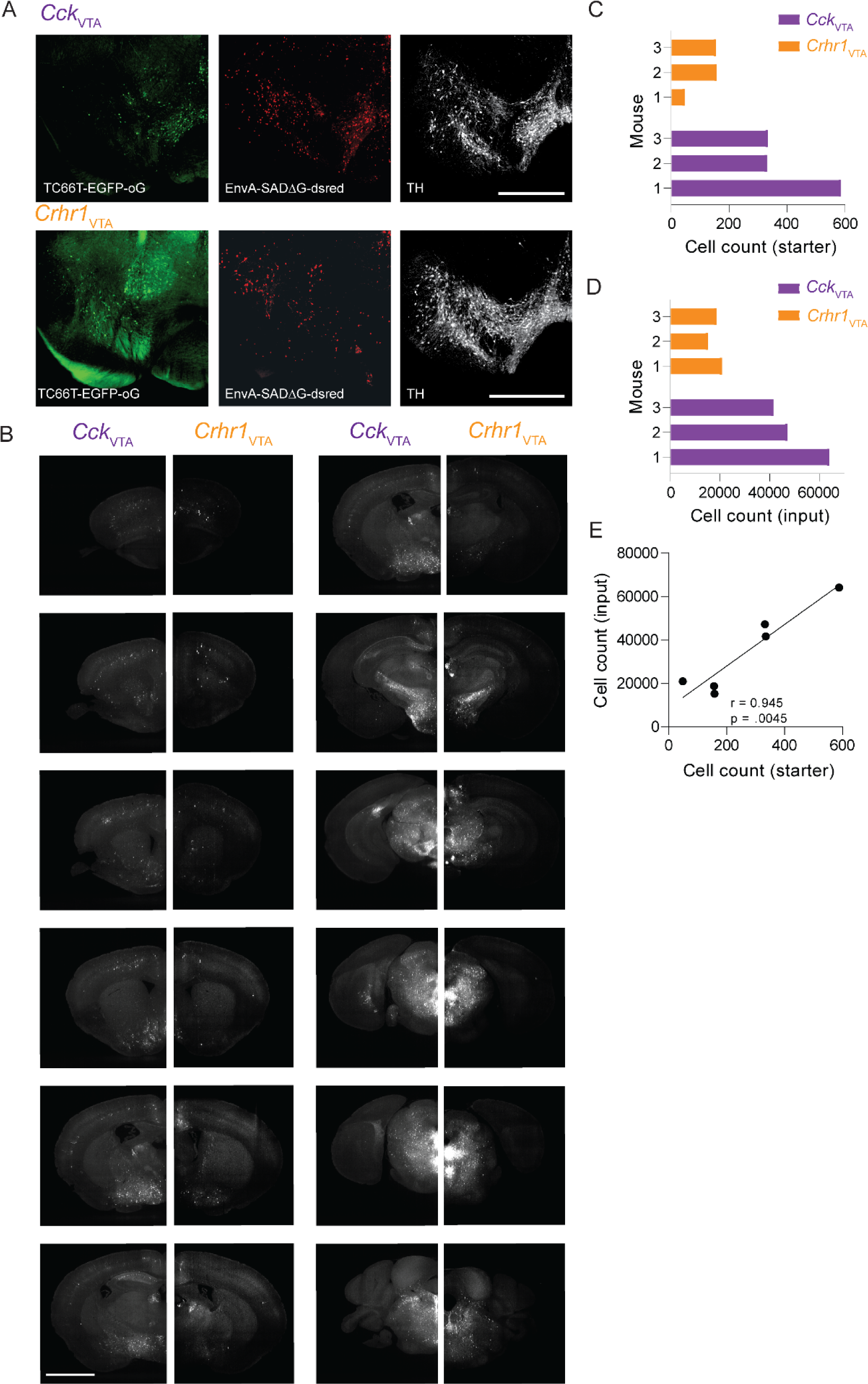
Further characterization of whole-brain input mapping to *Crhr1*_VTA_ and *Cck*_VTA_ neurons, related to Figure 8. (A) Representative images of *Cck*_VTA_ (top) and *Crhr1*_VTA_ (bottom) starter cell populations. Example histology images from the VTA showing staining for helper virus (AAV-syn-DIO-TC66T-2A-eGFP-2A-oG) (green), rabies virus (EnvA-SADΔG-RV-DsRed) (red), and tyrosine hydroxylase (white). Scale bar: 500 µm. (B) Representative images of *Cck*_VTA_ (left) and *Crhr1*_VTA_ (right) input cells. Scale bar: 2 mm. (C) Number of starter cells per mouse. (D) Number of labeled input neurons per mouse. (E) Relationship between numbers of starter and input neurons. Correlation coefficient (r) and p-value on the bottom right of the plot (see Supplementary Table 1 for statistical values).

**Figure S7.**
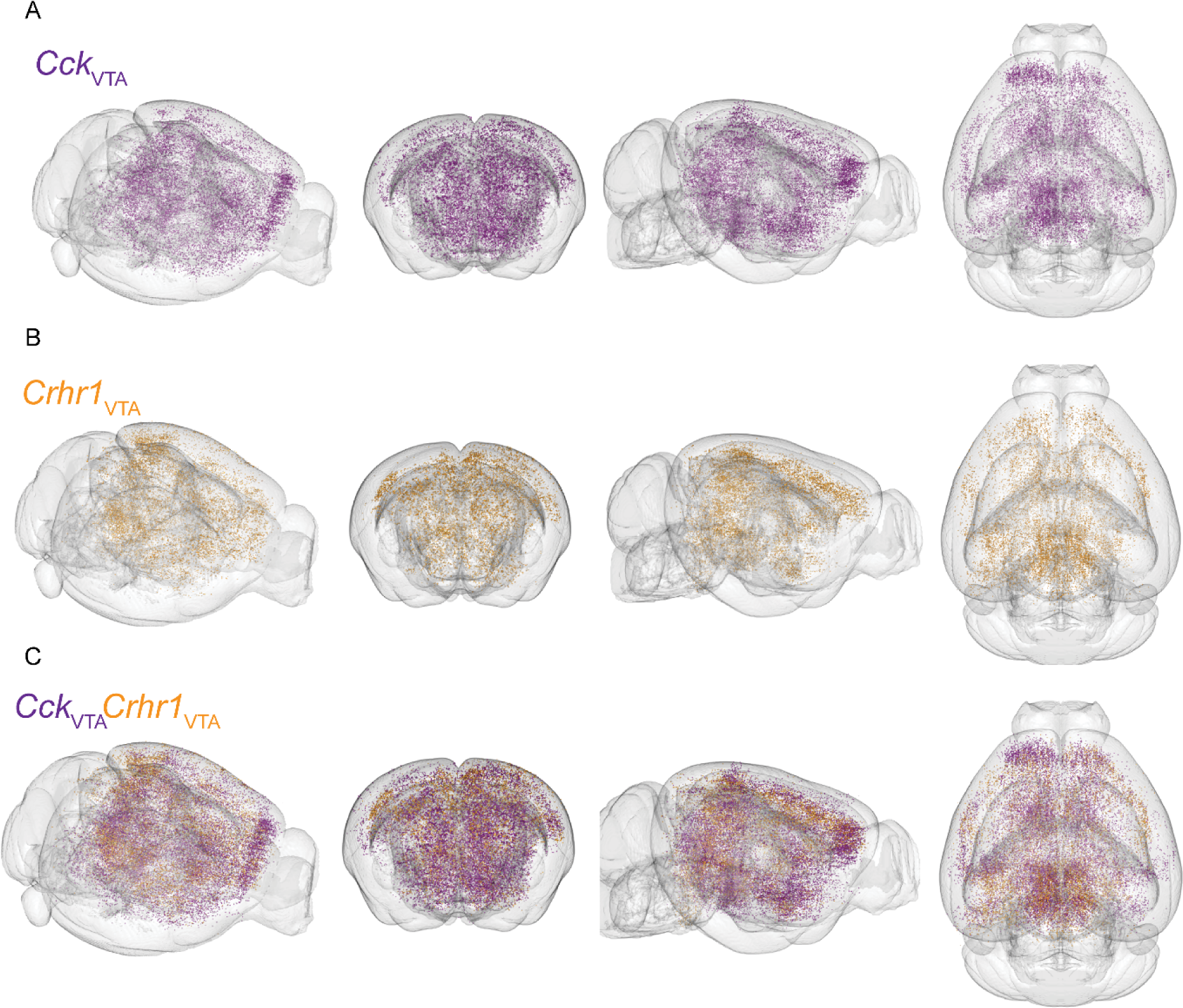
Further characterization of whole-brain input mapping to *Crhr1*_VTA_ and *Cck*_VTA_ neurons, related to Figure 8. (A) Location of input cells to *Cck*_VTA_ neurons in example mice. (B) Location of input cells to *Crhr1*_VTA_ neurons in example mice. (C) Location of input cells to *Cck*_VTA_ and *Crhr1*_VTA_ neurons in example mice.

**Supplementary Table 1.**
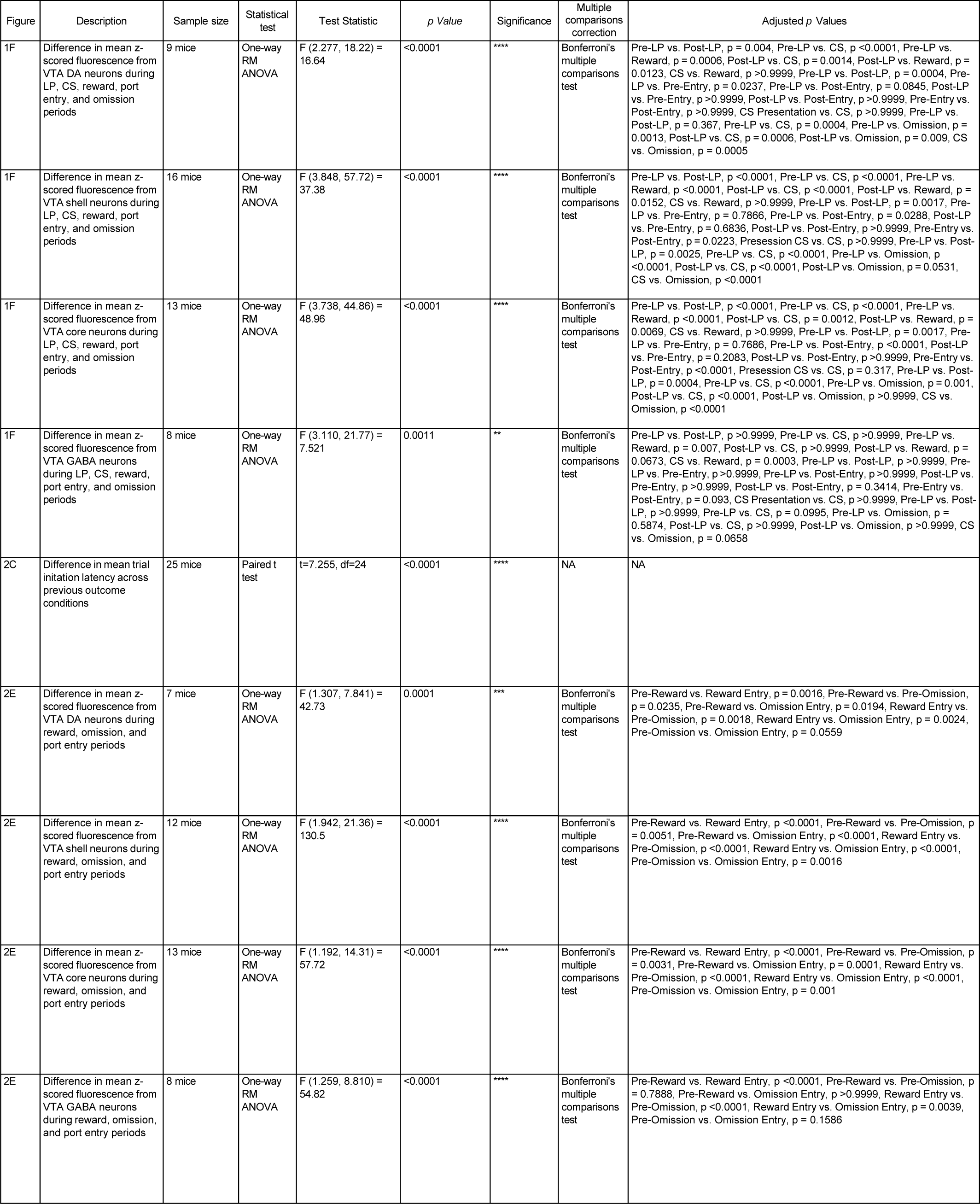
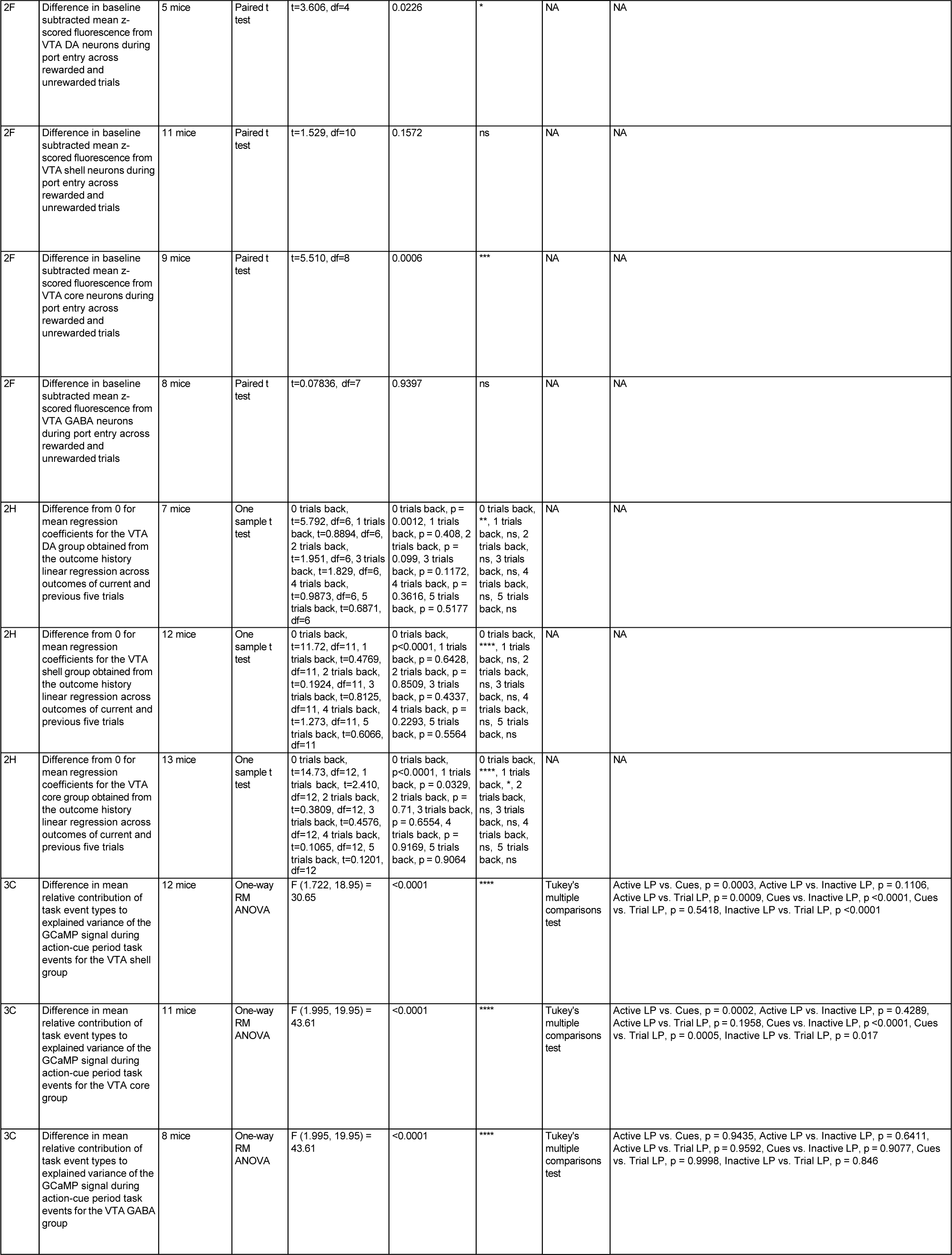
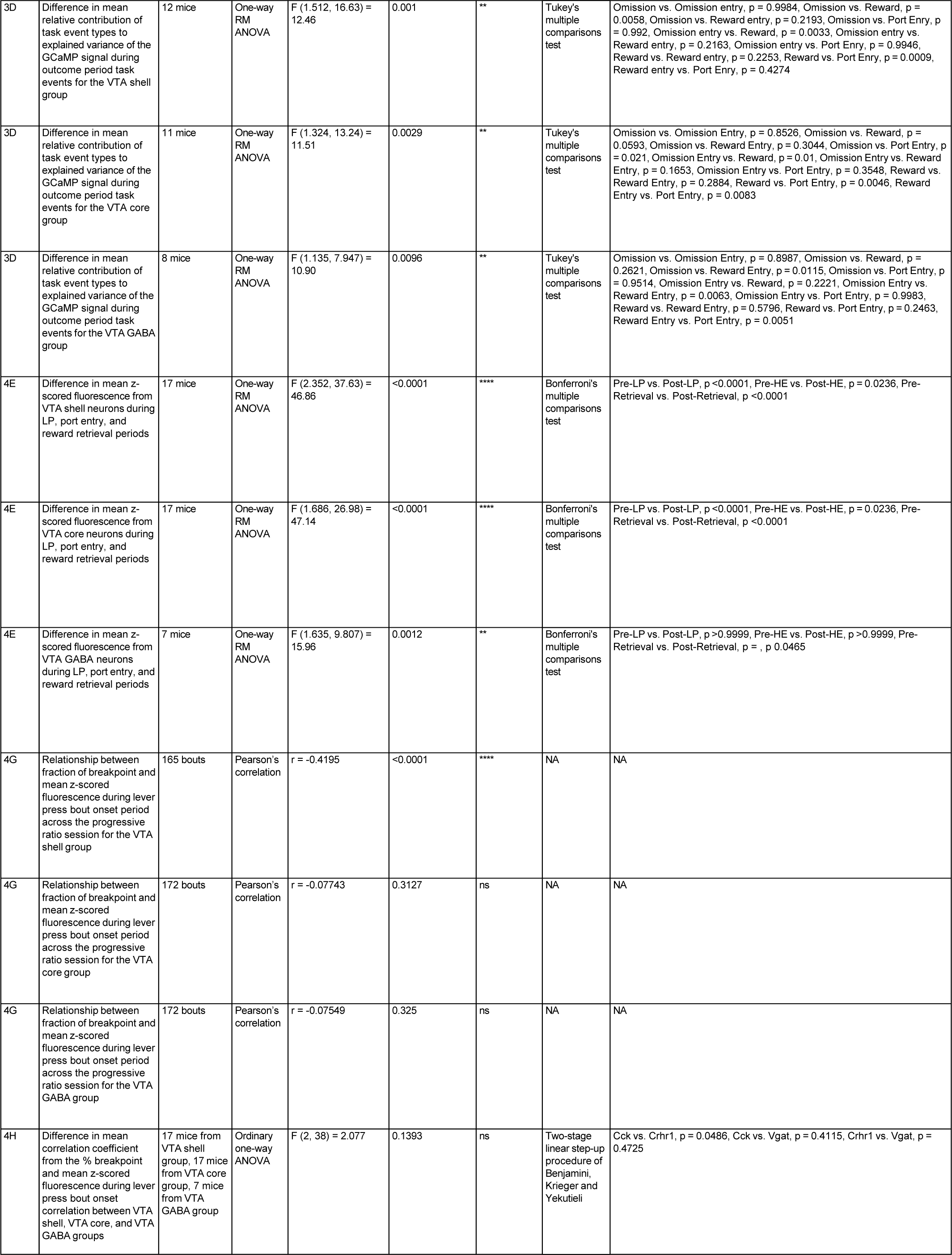
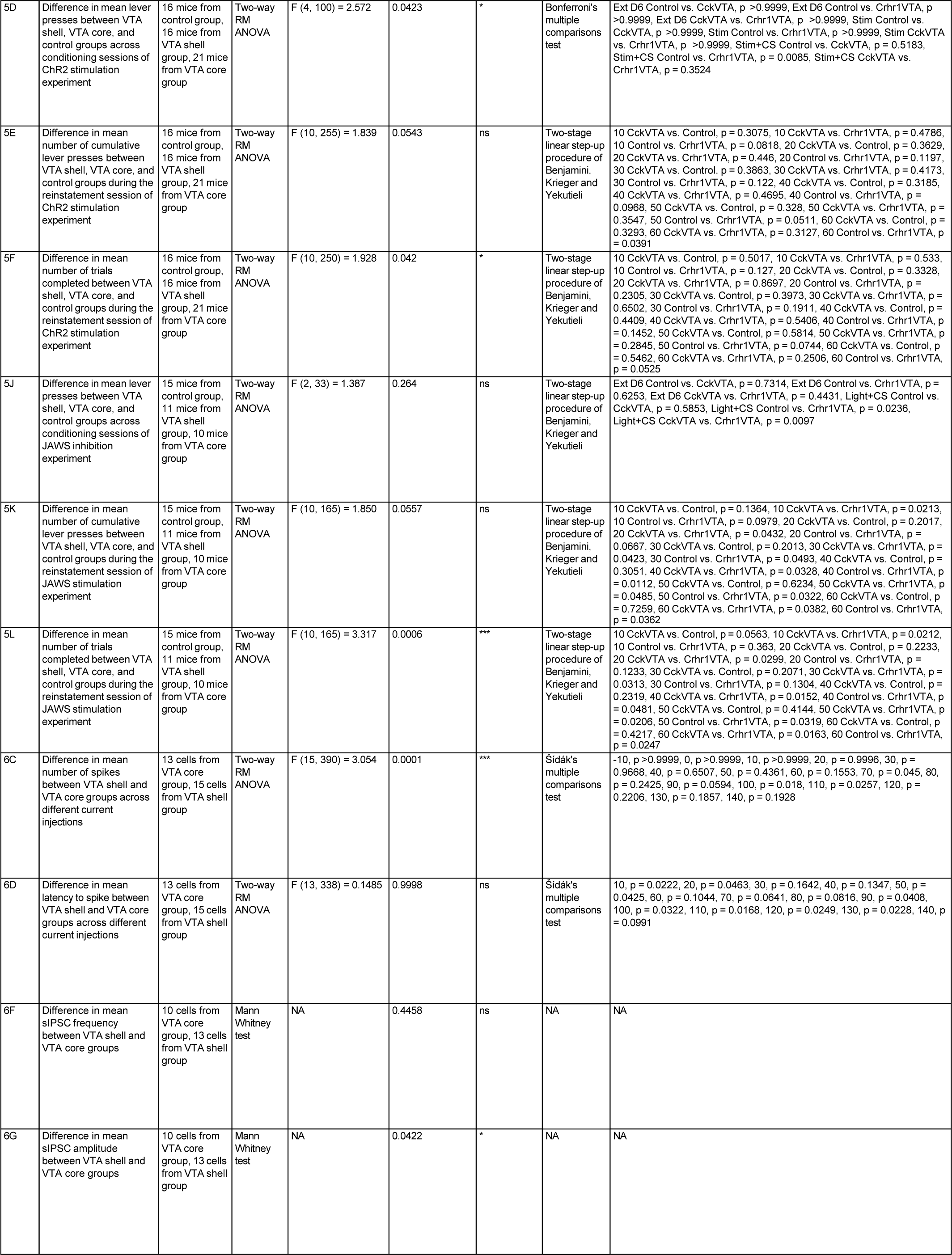
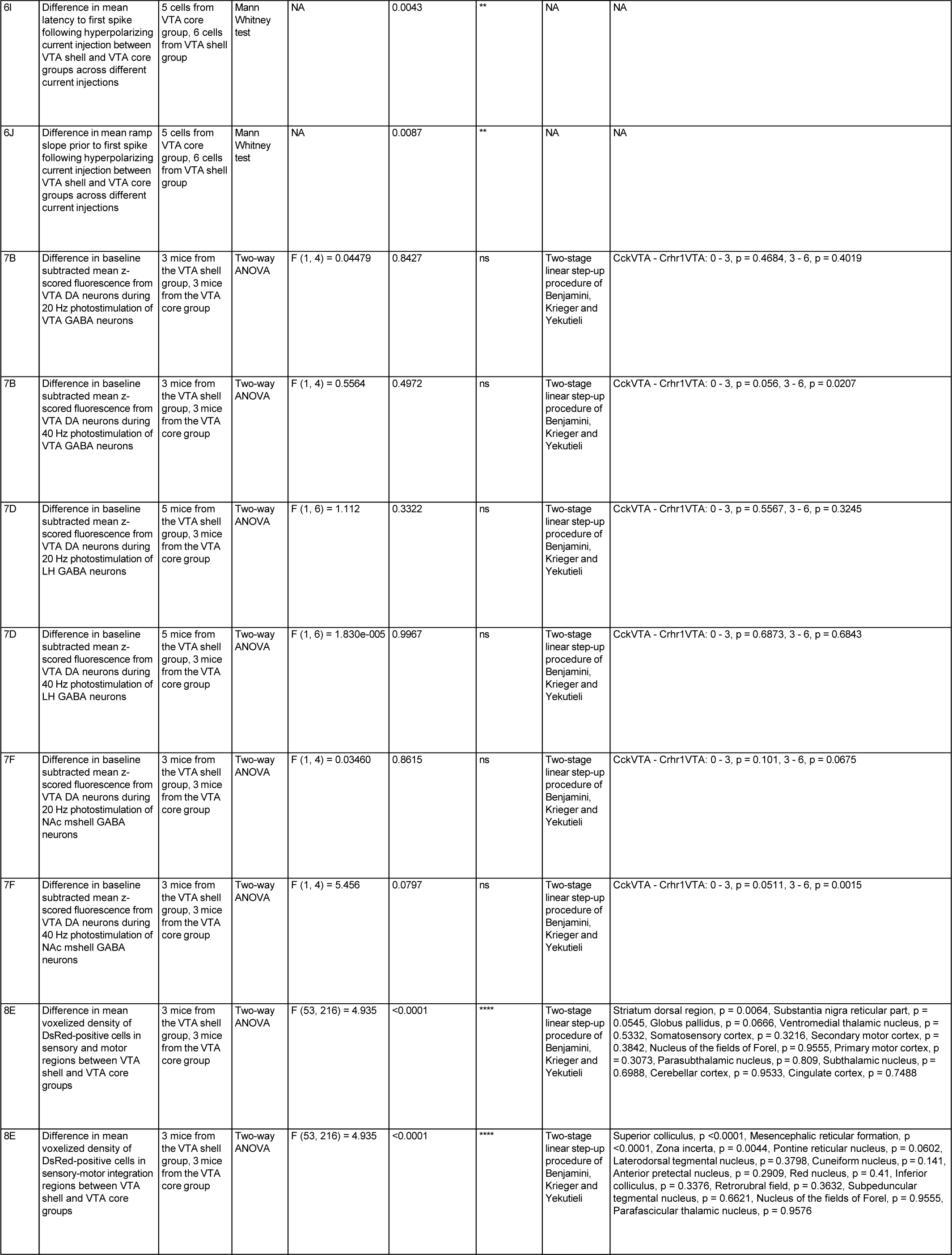
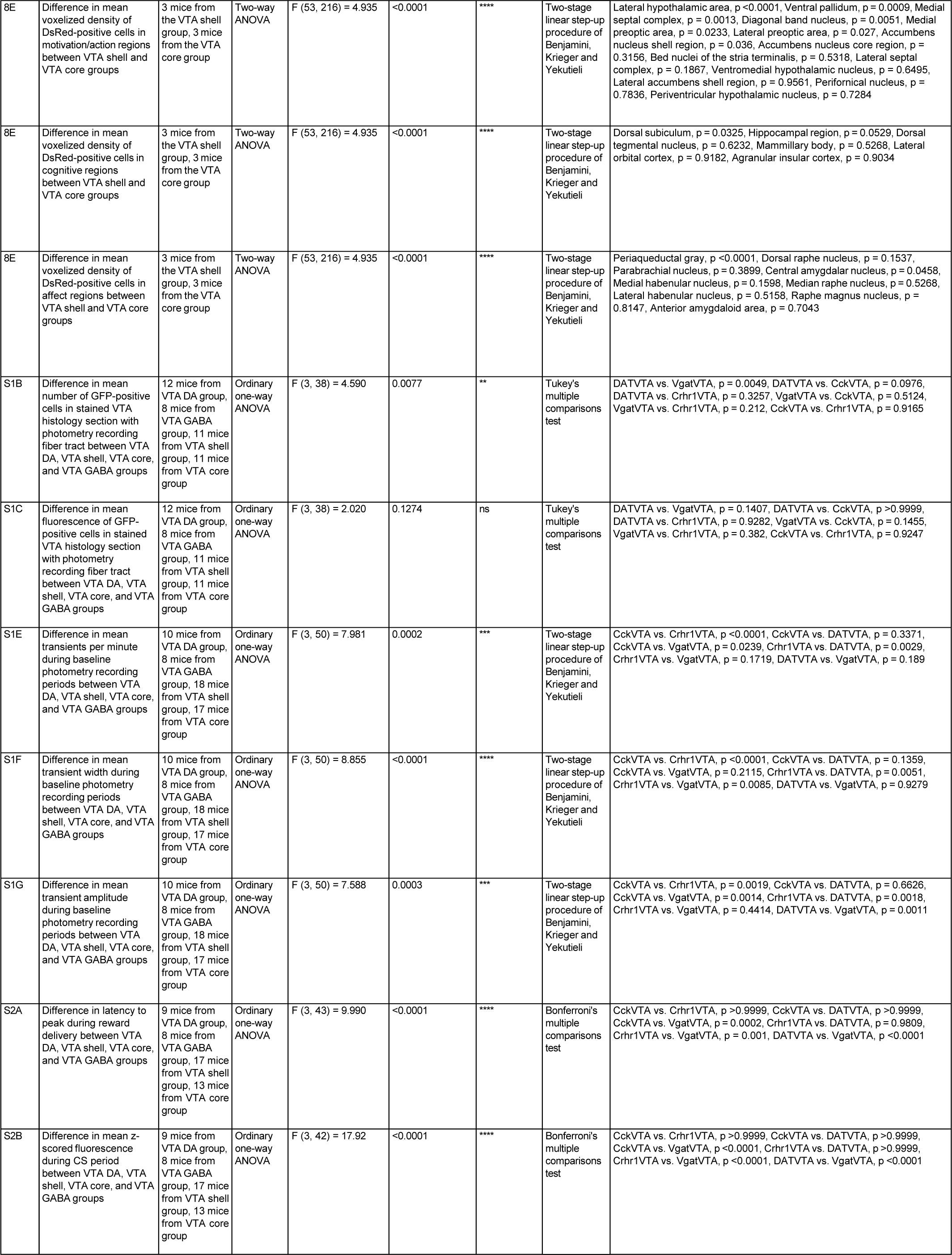
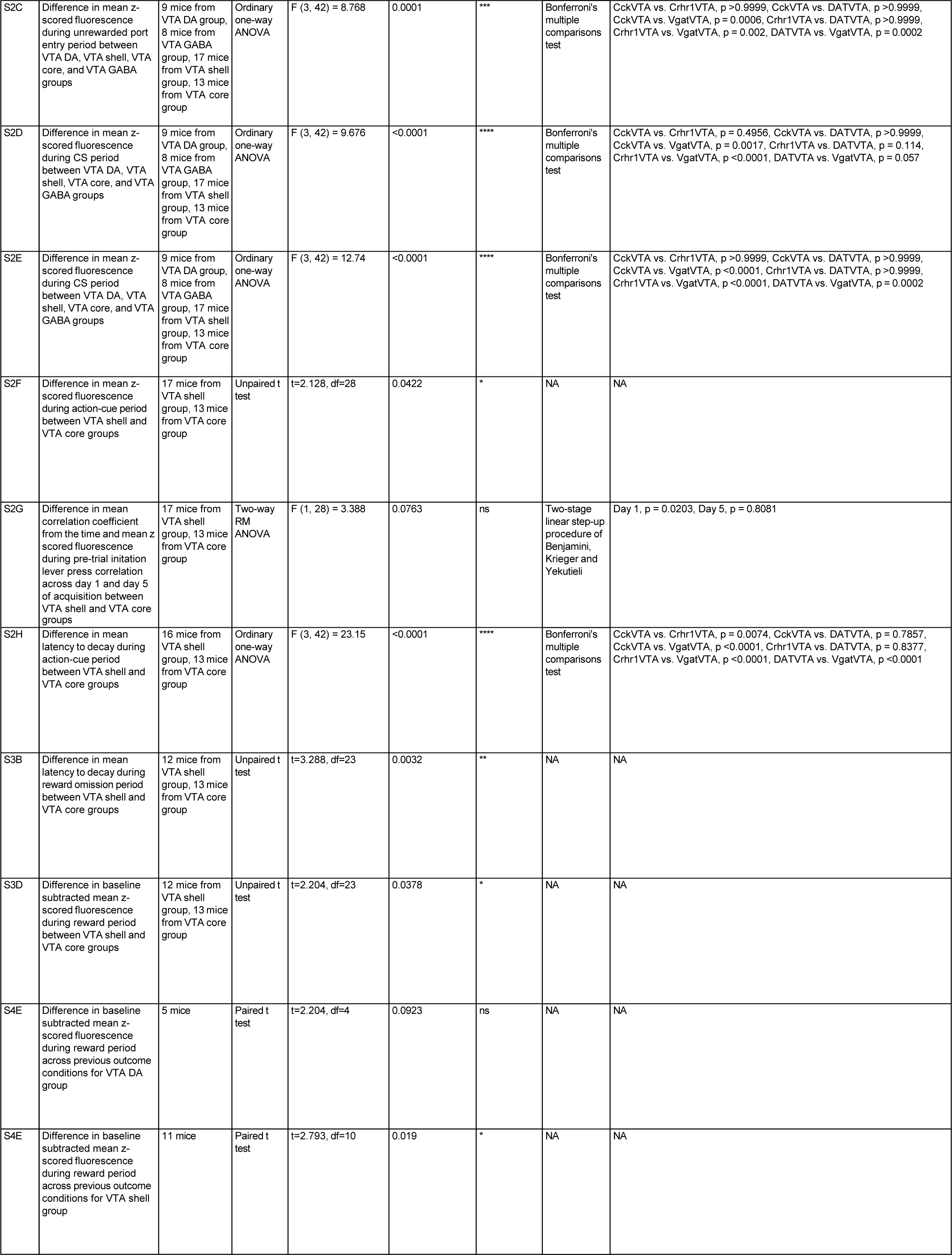
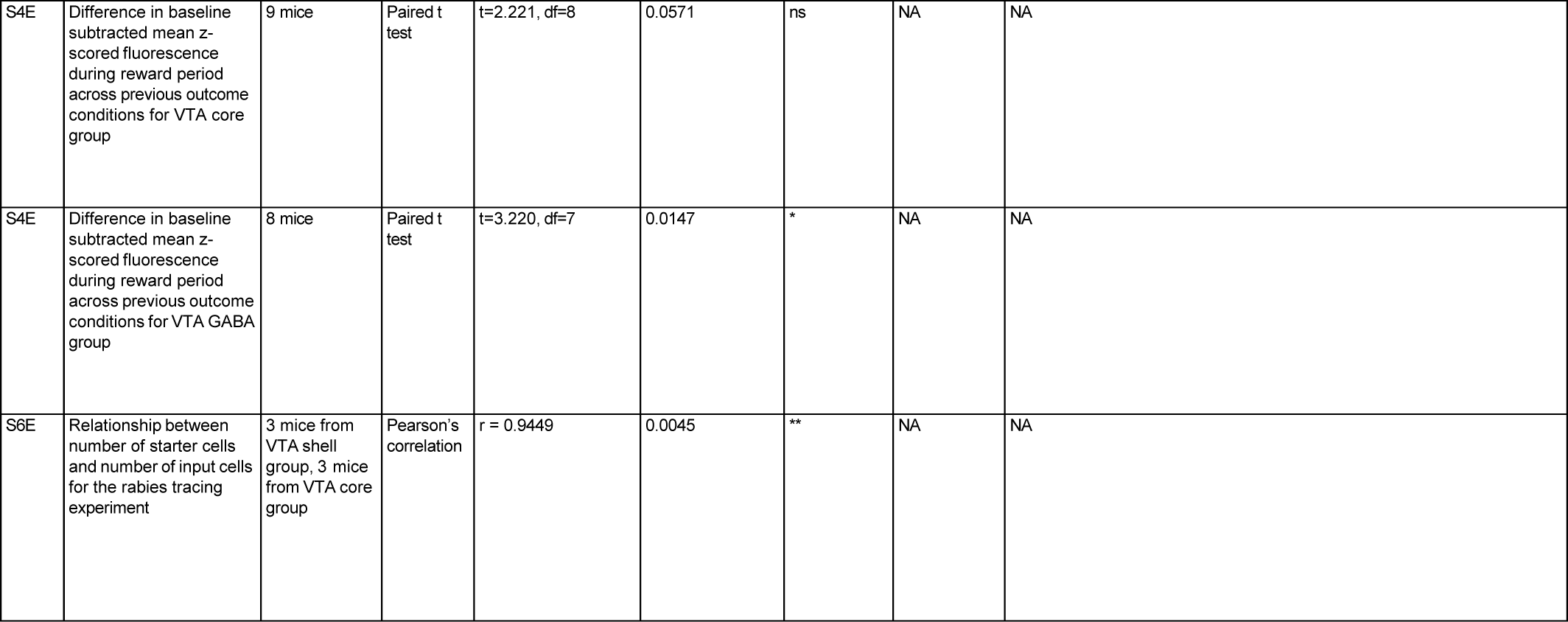
Statistical Results.

